# The CDK Pef1 and Protein Phosphatase 4 oppose each other for regulating cohesin binding to fission yeast chromosomes

**DOI:** 10.1101/748855

**Authors:** Adrien Birot, Marta Tormos-Pérez, Sabine Vaur, Amélie Feytout, Julien Jaegy, Dácil Alonso Gil, Stéphanie Vazquez, Karl Ekwall, Jean-Paul Javerzat

## Abstract

Cohesin has essential roles in chromosome structure, segregation and repair. Cohesin binding to chromosomes is catalyzed by the cohesin loader, Mis4 in fission yeast. How cells fine tune cohesin deposition is largely unknown. Here we provide evidence that Mis4 activity is regulated by phosphorylation of its cohesin substrate. A genetic screen for negative regulators of Mis4 yielded a CDK called Pef1, whose closest human homologue is CDK5. Inhibition of Pef1 kinase activity rescued cohesin loader deficiencies. In an otherwise wild-type background, Pef1 ablation stimulated cohesin binding to its regular sites along chromosomes while ablating Protein Phosphatase 4 had the opposite effect. Pef1 and PP4 control the phosphorylation state of the cohesin kleisin Rad21. The CDK phosphorylates Rad21 on Threonine 262. Pef1 ablation, non phosphorylatable Rad21-T262 or mutations within a Rad21 binding domain of Mis4 alleviated the effect of PP4 deficiency. Such a CDK/PP4 based regulation of cohesin loader activity could provide an efficient mechanism for translating cellular cues into a fast and accurate cohesin response.

## INTRODUCTION

Cohesin is a central player in chromosome biology. Defects in the cohesin pathway are linked to human pathologies such as sterility, cancer and severe developmental disorders [1–3]. Cohesin is an ATPase driven molecular machine able to tether DNA by topological entrapment [4–6]. The core cohesin complex is made of two long Structural Maintenance of Chromosome (SMC) proteins (Psm1 and Psm3 in fission yeast) whose ATPase heads are bridged by a kleisin subunit called Scc1/Mcd1/Rad21. The cohesin complex ensures proper chromosome segregation by holding sister chromatids together from DNA replication and until their segregation at anaphase onset. Besides chromosome segregation, cohesin is essential for DNA repair and the formation of DNA loops that shape chromosome architecture and impinge on gene regulation [7–9]. Cohesin is therefore central to many biological processes, emphasizing the importance of understanding its regulation. Cohesin loading onto DNA requires ATP hydrolysis by cohesin, which is thought to trigger transient opening of the ring and DNA entrapment [10–14]. The reaction is stimulated by a separate complex, the cohesin loader, made of Scc2-Scc4 in yeast, NIPBL-MAU2 in human, Mis4-Ssl3 in fission yeast [15–17]. Structural studies indicate that Scc2 consists of an N-terminal globular domain that binds Scc4 followed by helical repeats that fold into a hook-shaped structure [18; 19]. Mis4 alone has DNA binding activity and the C terminal hook domain is sufficient for stimulating DNA capture by cohesin *in vitro* [18; 20]. Mis4 makes multiple contacts with cohesin which may help cohesin conformational changes required for DNA capture [19; 20]. Although dispensable *in vitro*, Ssl3 is essential for cohesin binding to chromosomes [17; 20]. Scc4 wraps around Scc2 N terminus and is thought to recognize and bind chromatin receptors thereby directing cohesin loading to specific locations [18; 20–25]. Once loaded onto chromosomes cohesin can either remain bound or removed. Wpl1 promotes cohesin release in a reaction requiring Pds5 and Psc3. All three proteins bind Rad21 and Wpl1 promotes DNA release by weakening the Rad21-Smc3 interface [26–32]. DNA release is counteracted by a cohesin acetyl-transferase (Eso1 in fission yeast) that acetylates Smc3 during S phase [28; 33–36].

As opposed to sister-chromatid cohesion, cohesin loops may not require topological entrapment of DNA [14]. Cohesin may capture small loops of DNA and then extrude them in a processive manner. The formation of cohesin loops is dependent on the loading complex and reciprocally loops are de-stabilized by Wpl1 [37; 38]. Topological versus non topological DNA capture may be achieved by modulating the catalytic activity of the loader and concerted transcriptional responses may involve a local and temporal control of loading and unloading activities. How cells orchestrate cohesin functions is largely unknown. Intriguingly, the kleisin subunit of cohesin is targeted by multiple phosphorylation events. In fission yeast, Rad21 shows multiple phospho-isoforms whose relative abundance fluctuates along the cell cycle [39]. Our recent work showed that Protein Phosphatase 4 (PP4) controls the phosphorylation status of Rad21 and modulates Wpl1 activity [40], leading to the idea that cohesin functions could be spatially and temporally fine-tuned by altering the balance between kinase and phosphatase activities.

Here we report on the control of cohesin deposition by the opposite activities of the Pef1 CDK and PP4. Pef1 was first described as a PSTAIRE-related protein in fission yeast [41]. The CDK has three known cyclin partners called Pas1, Psl1 and Clg1 and was reported to facilitate the G1 to S phase transition and to regulate life span [42; 43]. Its closest human homolog, CDK5, is involved in a myriad of cellular functions and pathologies, from neurodegenerative diseases to multiple solid and hematological cancers [44]. We identified *pef1* in a genetic screen for mutants able to rescue the cohesin loader mutant *mis4-367*. Pef1 ablation or inhibition of its kinase activity increased cohesin deposition and rescued sister-chromatid cohesion defects of the *mis4* mutant. In otherwise wild-type cells Pef1 ablation increased the binding of both cohesin and its loader to their regular sites along chromosomes. Genetic analyses indicated that Pef1 acts through the phosphorylation of multiple targets. We identified one of these within the kleisin Rad21. Specifically, the Pef1/Psl1 complex phosphorylates Rad21 on T262 and preventing this phosphorylation event recapitulates in part the effects of Pef1 ablation. PP4 had the opposite effect. Its ablation lead to hyper-phosphorylated Rad21 and reduced cohesin deposition which is alleviated by Pef1 ablation or Rad21-T262A. Hence, phosphorylation of the kleisin negatively regulates cohesin loading, possibly by lowering the activity of the cohesin loader. Further supporting this notion, a genetic screen identified compensatory mutations that cluster within the catalytic domain of Mis4, in a previously described Rad21 binding region. Such a phosphorylation-based control may provide a fast, accurate and reversible way for regulating cohesin functions in response to cellular cues.

## RESULTS

### Inhibition of Pef1 kinase activity in *mis4-367* cells increases cohesin binding to DNA in S phase and improves chromosome segregation during mitosis

The *mis4-367* allele encodes Mis4^G1487D^. This single amino acid change is located within the last HEAT repeat of the C terminal catalytic domain (Fig. 1A), rendering the strain thermosensitive for growth (ts). To identify putative regulators of Mis4, we made a genetic screen for suppressors of the ts phenotype, the rationale being that loss of a negative regulator should upregulate residual Mis4^G1487D^ activity and restore growth at the restrictive temperature. Eleven mutants were isolated that distributed into 4 linkage groups. Genetic mapping and tiling array hybridization were used to identify the mutated locus in group 1. A single base substitution was found within the *pef1* coding sequence. The amino acid change (N146S) is located within the catalytic site of the kinase suggesting the kinase activity was involved. Accordingly, deletion of the *pef1* gene or inhibition of Pef1 kinase activity using an analog sensitive allele (*pef1-as*) suppressed the ts growth defect of *mis4-367* (Fig. 1B).

**Figure 1.**
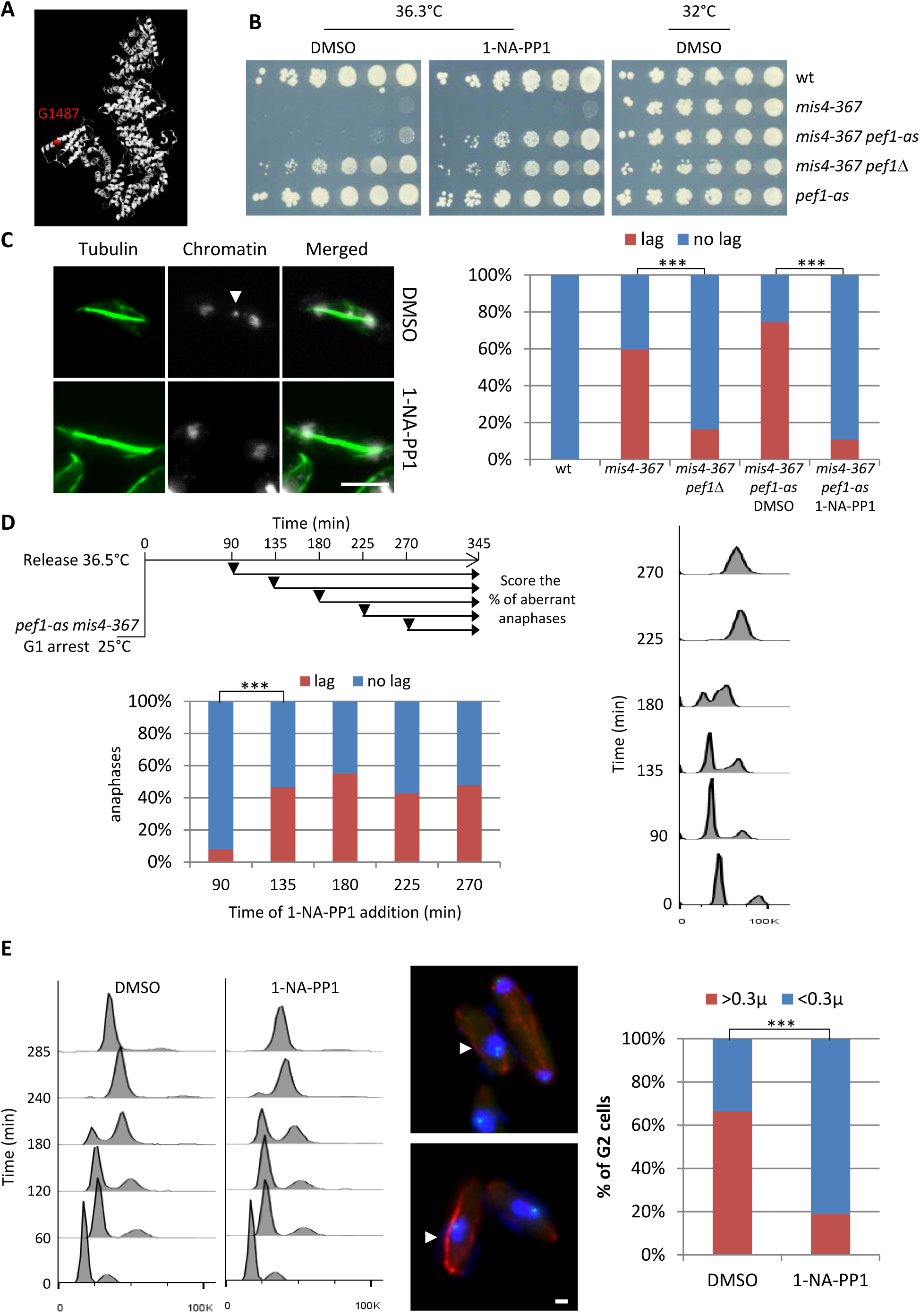
Inhibition of Pef1 kinase activity suppressed Mis4^G1487D^ cohesion and chromosome segregation defects. **(A)** The *mis4-367* allele results in a G1487D substitution within the last HEAT repeat of Mis4. **(B)** Cell growth assay showing that inhibition of Pef1 kinase activity suppresses *mis4-367* thermosensitive growth phenotype. **(C)** Inhibition of Pef1 kinase activity suppresses *mis4-367* chromosome segregation defects. Cells were cultured at 36.5°C for a complete cell cycle. Lagging chromatids appear as DAPI-stained material (arrow) along the anaphase spindle (tubulin staining in green). Bar = 5µm. ***p< 0.0001 two-sided Fisher’s exact test. **(D)** Pef1 inhibition must occur before S phase onset to rescue *mis4-367* chromosome segregation. Cells were arrested in G1 by nitrogen starvation, released into the cell cycle at 36.5°C and 1-NA-PP1 added at the indicated time points (arrows). Cell cycle progression was followed by measurement of DNA content. Anaphase cells with lagging chromatids were scored at the 345min time point. ***p< 0.0001 two-sided Fisher’s exact test. **(E)** Pef1 kinase inhibition improved sister-chromatid cohesion. Cells were arrested in G1 by nitrogen starvation, released into the cell cycle at 36.5°C with or without 1-NA-PP1. Cells were harvested after DNA replication (285min) and processed for FISH using a centromere 2-linked probe. Distance between FISH signals was measured in G2 cells, as judged by DNA content and the interphase array of microtubules. Bar = 1µm. ***p< 0.0001 two-sided Fisher’s exact test.

Likewise, *pef1*Δ showed a suppressor effect towards the strong ts allele *mis4-242* [45] and efficiently suppressed *ssl3-29* (Fig. S1), a ts mutant of *ssl3* [17]. The deletion of *pef1* even allowed cell survival in the complete absence of the *ssl3* gene, although colonies were tiny and grew very slowly (Fig. S1). By contrast *pef1*Δ showed a negative genetic interaction with *eso1-H17* [46] indicating that *pef1* displays distinct genetic interactions with components of the cohesin pathway (Fig. S1). Deletion of *pef1* did not allow cell survival in the complete absence of the *mis4* gene (Fig. S1), indicating that *pef1* mediated suppression required residual Mis4 activity. Altogether, these genetic data suggested that *pef1* deletion may upregulate Mis4. The corollary being that the CDK may act as a negative regulator of Mis4. We first aimed at characterizing the suppression of Mis4^G1487D^ phenotypes by *pef1* mutants. Thermosensitive mutants of the cohesin loading complex fail to properly establish sister-chromatid cohesion during S phase [15; 17; 47] and consequently display a high frequency of aberrant mitoses in which sister chromatids lag along the spindle during anaphase. After one complete cell cycle at the restrictive temperature *mis4-367* cells displayed a high frequency of aberrant anaphases, a defect which was efficiently rescued by the deletion of the *pef1* gene. The chemical inhibition of Pef1-as had a similar effect, confirming that Pef1 acts through its kinase activity (Fig. 1C).

The cohesin loading complex performs its essential function during G1/S [15; 17; 47]. Accordingly chromosome segregation was efficiently restored at 36.5°C when Pef1-as was inhibited before but not after S phase onset (Fig. 1D). Finally sister-chromatid cohesion was monitored by FISH, using a probe located close to the centromere of chromosome 2 (Fig. 1E). The inhibition of Pef1 kinase activity significantly reduced the frequency of separated FISH signals, consistent with improved sister chromatid cohesion.

The failure to establish cohesion during S phase may be due to poor cohesin loading. To see whether the inhibition of Pef1 kinase activity would improve cohesin binding to chromosomes at the time of cohesion establishment, *mis4-367 pef1-as* cells were cultured in the presence of 1-NA-PP1 or solvent alone (DMSO) and arrested in S phase at the restrictive temperature (Fig. 2A). The amount of chromatin bound Rad21 was monitored by Chromatin Immunoprecipitation (ChIP) at known Cohesin Associated Regions (CARs) along the arms and centromere of chromosome 2, the rDNA gene cluster on chromosome 3 and the chromosome 1 right telomere (Fig. 2B) [40; 48]. Within the centromere, Rad21 binding was examined at the central core (cc2) which is the site of kinetochore assembly, within the *imr* and *dg* repeats that flank the central core on either side and at tRNA rich domains that delineate the centromere. The peri-centromere repeats are strong cohesin binding sites in wild-type and provide robust sister chromatid cohesion at centromeres [49; 50]. As expected Rad21 binding was reduced in the *mis4-367* mutant as compared to wild-type (Fig. 2C). The ratio 1-NA-PP1 / DMSO indicated that Rad21 binding was increased in the presence of 1-NA-PP1 in a *pef1-as* dependent manner (Fig. 2D). The ratios in Fig. 2E indicate that Rad21 binding was restored to ∼60% the wild-type level within the rDNA Non Transcribed Spacer (NTS) and ∼50% at the telomere site. Within the centromere, Rad21 binding was back to ∼50% wild-type levels within the outer repeats (*imr2-L* and *dg2-R*), consistent with improved sister-chromatid cohesion as seen with the cen2 FISH assay (Fig. 1E).

**Figure 2.**
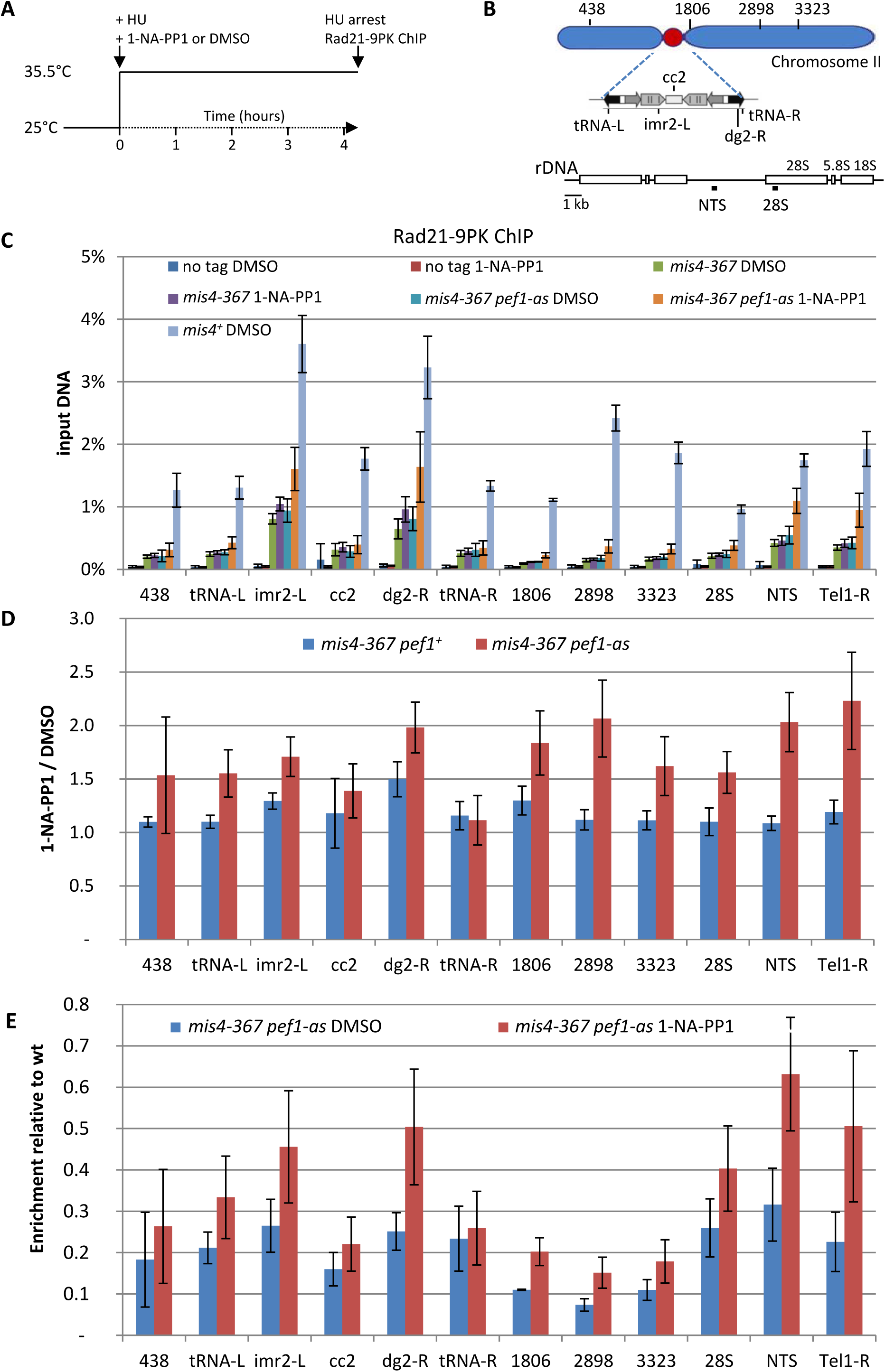
Inhibition of the CDK Pef1 in *mis4-367* increased Rad21 binding to S phase chromosomes. **(A)** Scheme of the experiment. Hydroxyurea (HU) was added to 12mM at the time of the temperature shift along with 1-NA-PP1 or solvent alone (DMSO). Cells were collected after 4.25 hours. **(B)** Schematics showing the loci analyzed by ChIP-qPCR. **(C)** Rad21-9PK ChIP-qPCR results expressed as % of input DNA. The “no tag” control estimates background enrichment. Bars indicate mean ± SD from 4 ChIPs. **(D)** The effect of 1-NA-PP1 treatment is shown by the ratio 1-NA-PP1/DMSO (red) for each site analyzed. The ratios in a *pef1^+^* background (blue) estimate the off target effects of the inhibitor. Ratios were calculated from the data shown in (C). Bars indicate mean ± SD from 4 ratios. **(E)** Rad21 binding relative to wild-type. The ratios highlight the recovery of Rad21 binding upon inhibition of the CDK relative to wild-type levels. Ratios were calculated from the data shown in (C). Bars indicate mean ± SD from 4 ratios.

The total amount of chromatin bound Rad21 per nucleus, as assayed by nuclear spreads (Fig. S2), remained largely unchanged. This may indicate that the increase in Rad21 binding is mainly confined to CARs and the resulting global increase is below the sensitivity of this assay.

The establishment of sister chromatid cohesion in S phase is accompanied by acetylation of the cohesin subunit Psm3 [35]. At 25°C Psm3 K106 acetylation in *mis4-367* cycling cells was similar to wild-type (Fig. S2). By contrast, the level dropped in S phase arrested *mis4-367* cells at 35.5°C. Psm3 K106 acetylation was slightly increased when Pef1-as was inhibited consistent with enhanced cohesin binding to DNA, improved sister-chromatid cohesion establishment or both.

From this set of experiments we conclude that the inhibition of Pef1 kinase activity stimulates cohesin binding to CARs in Mis4 deficient cells and improves the establishment of sister chromatid cohesion and chromosome segregation.

In fission yeast, a small fraction of cohesin may dissociate from chromatin during early mitosis, another is cleaved by separase at anaphase onset while the bulk may remain bound to chromosomes [48; 51]. Pef1 inhibition may rescue *mis4-367* by acting on cohesin from the previous cell cycle and/or may stimulate *de novo* cohesin loading. The latter possibility was investigated by inducing the expression of an ectopic FLAG-tagged *rad21* construct in G1 arrested cells. Growth assays indicated that ectopically expressed *rad21-FLAG* was functional as it allowed cell division in the absence of the endogenous *rad21* gene and *pef1*Δ suppressed the *mis4-367* ts growth defect under these conditions (Fig. 3A). Rad21-FLAG binding to chromatin was monitored by cell fractionation (Fig. 3D). Neo-synthesized Rad21-FLAG was poorly associated with the chromatin fraction when *pef1-as* was not inhibited. By contrast, chromatin bound Rad21-FLAG was increased upon inhibition of the CDK, consistent with increased *de novo* cohesin loading.

**Figure 3.**
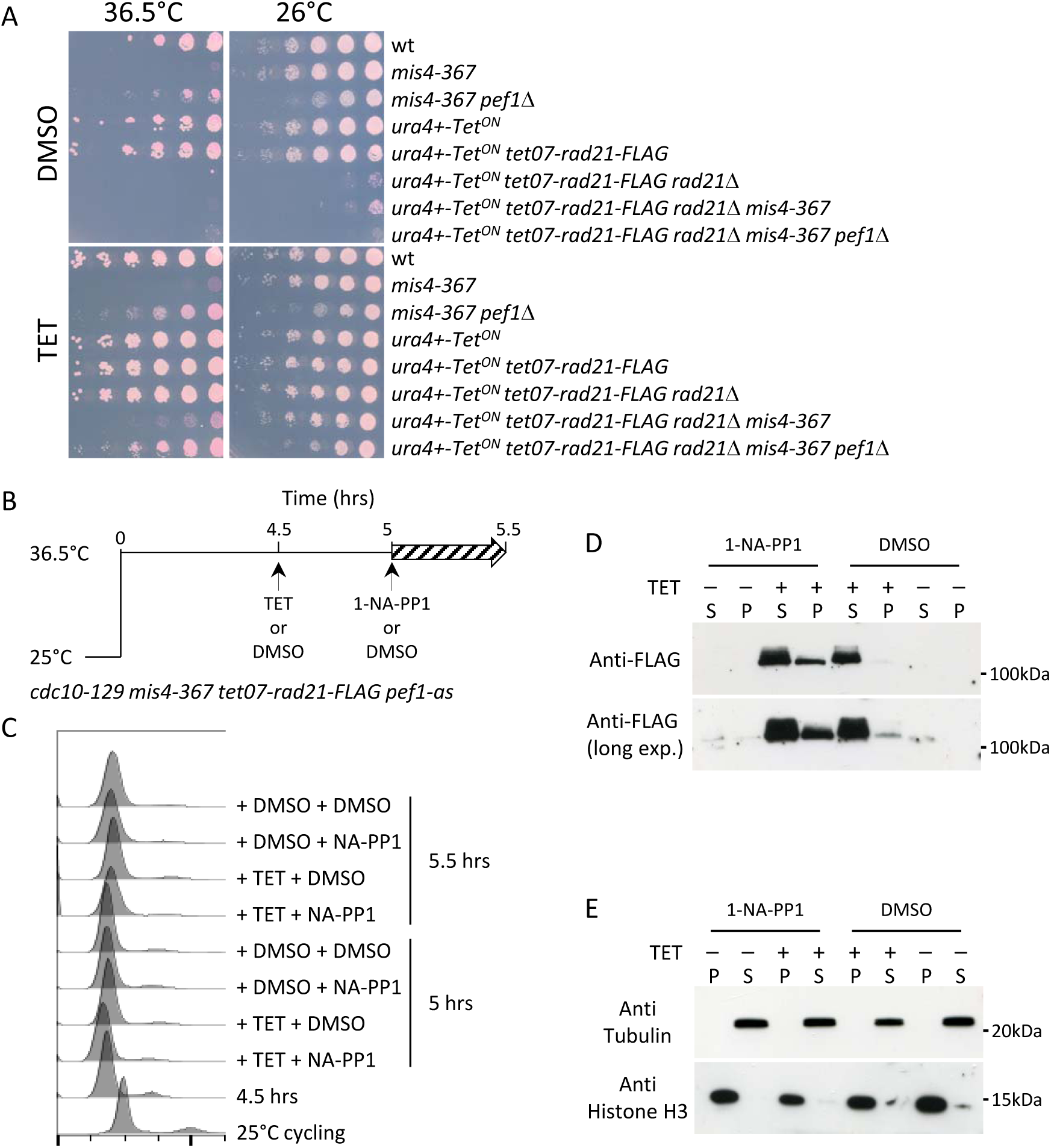
Inhibition of Pef1 kinase in G1 arrested *mis4-367* cells allows neo-synthesized Rad21 to bind chromatin. **(A)** The tetracycline (TET) inducible *tet07-rad21-FLAG* construct can substitute for the endogenous *rad21* gene. The last two lanes show that *pef1Δ* suppresses *mis4-367* ts phenotype when *tet07-rad21-FLAG* is the sole source of Rad21. **(B)** Experimental scheme. Cells cultured in EMM2 medium were arrested in G1 by the *cdc10-129* mutation. After 4.5hrs *tet07-rad21-FLAG* was induced by the addition of TET or left un-induced (DMSO). Pef1-as was inhibited 30min later and samples collected after 30min. **(C)** DNA content analysis. **(D)** Western blot analysis of Rad21-FLAG in the chromatin (P) and soluble (S) fractions. **(E)** Fractionation controls. Anti-tubulin and anti-Histone H3 antibodies were used as markers for the soluble (S) and chromatin (P) fractions, respectively.

Enhanced cohesin binding to DNA may result from increased cohesin loading, reduced unloading or both. Two mechanisms have been described for unloading cohesin from G1 chromosomes, both involving the opening of the Smc3/Kleisin interface. One is dependent on Wpl1 while the other is not and was recently reported in budding yeast to be inhibited by Scc2 [52]. If *pef1*Δ suppressed *mis4-367* by inhibiting such an unloading mechanism, the artificial closure of the Psm3/Rad21 interface should suppress *mis4-367*. As originally reported in budding yeast [30], a *psm3-rad21* gene fusion efficiently bypassed the requirement for the Eso1 acetyl-transferase (Fig. S3A) but did not restore and even enhanced the temperature growth defect of *mis4-367* (Fig. S3B). Importantly *pef1*Δ still showed a suppressor activity in this genetic setup, indicating that Pef1 acts independently from the Psm3/Rad21 interface. Likewise the deletion of *wpl1* had little effect, if any (Fig. S3C). These genetic data argue against a Pef1 mediated control of the Smc3/Rad21 interface. We therefore favor the conclusion that Pef1 inhibition rescued cohesin loader deficiency by increasing its residual activity.

### Chromatin binding of cohesin and its loader Mis4 are negatively regulated by Pef1

The above data suggest that the residual cohesin loading activity in *mis4-367* is enhanced when Pef1 kinase is inhibited, implying that the CDK may function as a negative regulator of Mis4. We addressed this question by looking at the effect of Pef1 ablation on Rad21 and Mis4 binding to chromosomes in otherwise wild-type cells. In cells arrested in G1 by the *cdc10-129* mutation, both Mis4 and Rad21 binding were increased in a correlated manner in *pef1*Δ cells (Fig. 4BCD). The strongest increase was observed at the centromere, telomere and NTS sites. Chromosome arm sites did respond as well although to a lesser extent. In cycling cells which are mainly (80%) in G2 [53], Rad21 and Mis4 binding were still enhanced within the centromere, NTS and telomere but barely changed along chromosome arm sites (Fig. 4E). Mis4 and Rad21 binding were also less well correlated, in contrast to G1 (Fig. 4D). A co-immunoprecipitation assay indicated that the amount of cohesin bound to Mis4 in G1-arrested cells was slightly enhanced in the absence of Pef1 (Fig. 4F). The effect was weak but consistent between experiments. This may reflect the increased abundance of both cohesin and its loader at CARs. It is of note that Rad21 was hypo-phosphorylated in the absence of the CDK (as detailed below) which may modify how cohesin, Mis4 and DNA interact with each other.

**Figure 4.**
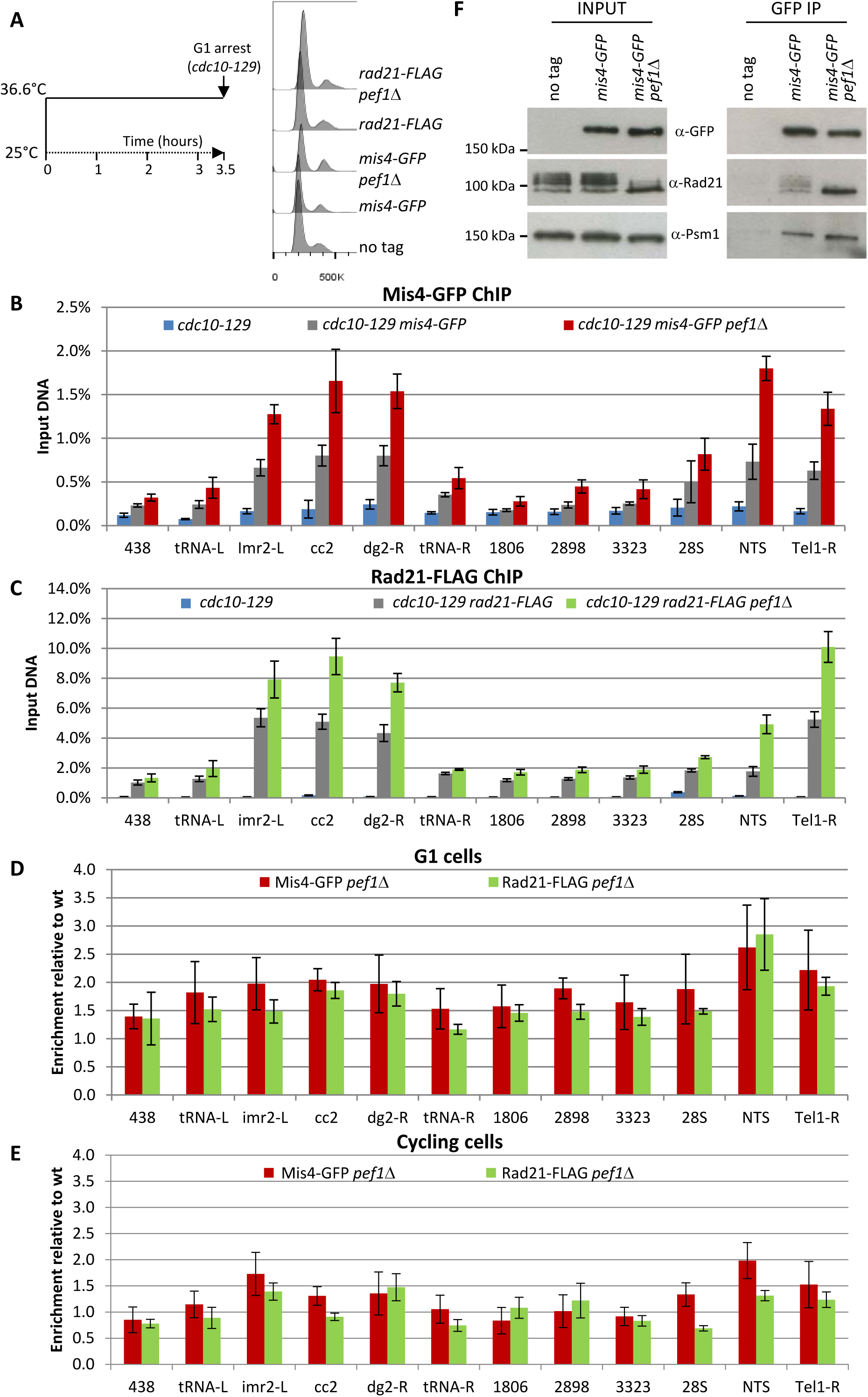
Pef1 ablation stimulates Rad21 and Mis4 binding to G1 chromosomes. **(A)** Strains of the indicated genotypes were arrested in G1 by the *cdc10-129* mutation. The G1 arrest was monitored by DNA-content analysis. Cells were processed for Mis4-GFP **(B)** and Rad21-FLAG **(C)** ChIP-qPCR at the indicated loci. Bars indicate mean ± SD from 4 ChIP. **(D)** Data from (B) and (C) were normalized to wild-type levels. Bars indicate mean ± SD from 4 ratios. **(E)** Rad21-FLAG and Mis4-GFP binding in cycling cells. ChIP data were normalized to wild-type levels. Bars indicate mean ± SD from 4 ratios. **(F)** Pef1 ablation increased cohesin binding to its loader. Mis4-GFP was immuno-purified from G1 (*cdc10-129*) protein extracts and co-purifying proteins were analyzed by western blotting with the indicated antibodies.

### The Pef1 CDK phosphorylates Rad21

Pef1 may act through the phosphorylation of one or several critical substrates. Pef1 co-immunoprecipitated cohesin and Mis4 from total protein extracts (Fig.5 AB) and western blot analyses indicated that the phosphorylation state of Rad21 was altered in *pef1* deleted cells (Fig. 5C). In wild-type, Rad21 displays multiple phospho-isoforms with reduced electrophoretic mobility by SDS-PAGE ([39; 40; 54] and Fig.5C). Fast migrating Rad21 species accumulated in *pef1* deleted cells at the expense of slow migrating forms. To see whether Pef1 directly phosphorylated Rad21 we used *in vitro* kinase assays. Indeed, the CDK purified from cycling or G1-arrested cells phosphorylated Rad21 (Fig. 5D). Pef1 was reported to bind three different cyclins. *In vitro* Rad21 phosphorylation was abolished when Pef1 was purified from *psl1* deleted cells (Fig. 5D), strongly suggesting that Pef1 acts together with the Psl1 cyclin to phosphorylate Rad21. Accordingly, the electrophoretic mobility of Rad21 was similar in *psl1*Δ and *pef1*Δ cells (Fig. 5C). To confirm that Pef1 uses the Psl1 cyclin to phosphorylate Rad21, *pef1-GFP* was fused to the endogenous *psl1* gene in a *pef1*Δ background so that Psl1-Pef1-GFP should be the sole Pef1 CDK available in the cell. Indeed the fusion protein purified from cell extracts phosphorylated Rad21 *in vitro* (Fig. 5E). In addition, the cohesin core subunit Psm1 efficiently co-purified with Psl1-GFP (Fig. 5F). To identify the phosphorylated residue(s), truncated Rad21 peptides were used as substrates for *in vitro* kinase assays (Fig. S5). A N-terminal fragment (1-356) was efficiently phosphorylated by Pef1 *in vitro*. Several CDK consensus sites (S/T-P) lie within that region. Replacement of T262 by an alanine abolished *in vitro* Rad21 phosphorylation by Pef1-GFP (Fig. 5E and S5). A weak signal was sometime observed with long exposure times (Fig. S5) or with the Psl1-Pef1 fusion protein (Fig. 5E), suggesting that other Rad21 residues might be additional or alternative substrates of the kinase. Finally, we raised antibodies against a Rad21-T262 phosphorylated peptide. As shown in Fig. 5G, the antibodies detected Rad21 purified from wild-type but not from *pef1*Δ cells extracts. From this set of experiment we conclude that the Pef1/Psl1 CDK phosphorylates Rad21 on threonine 262.

**Figure 5.**
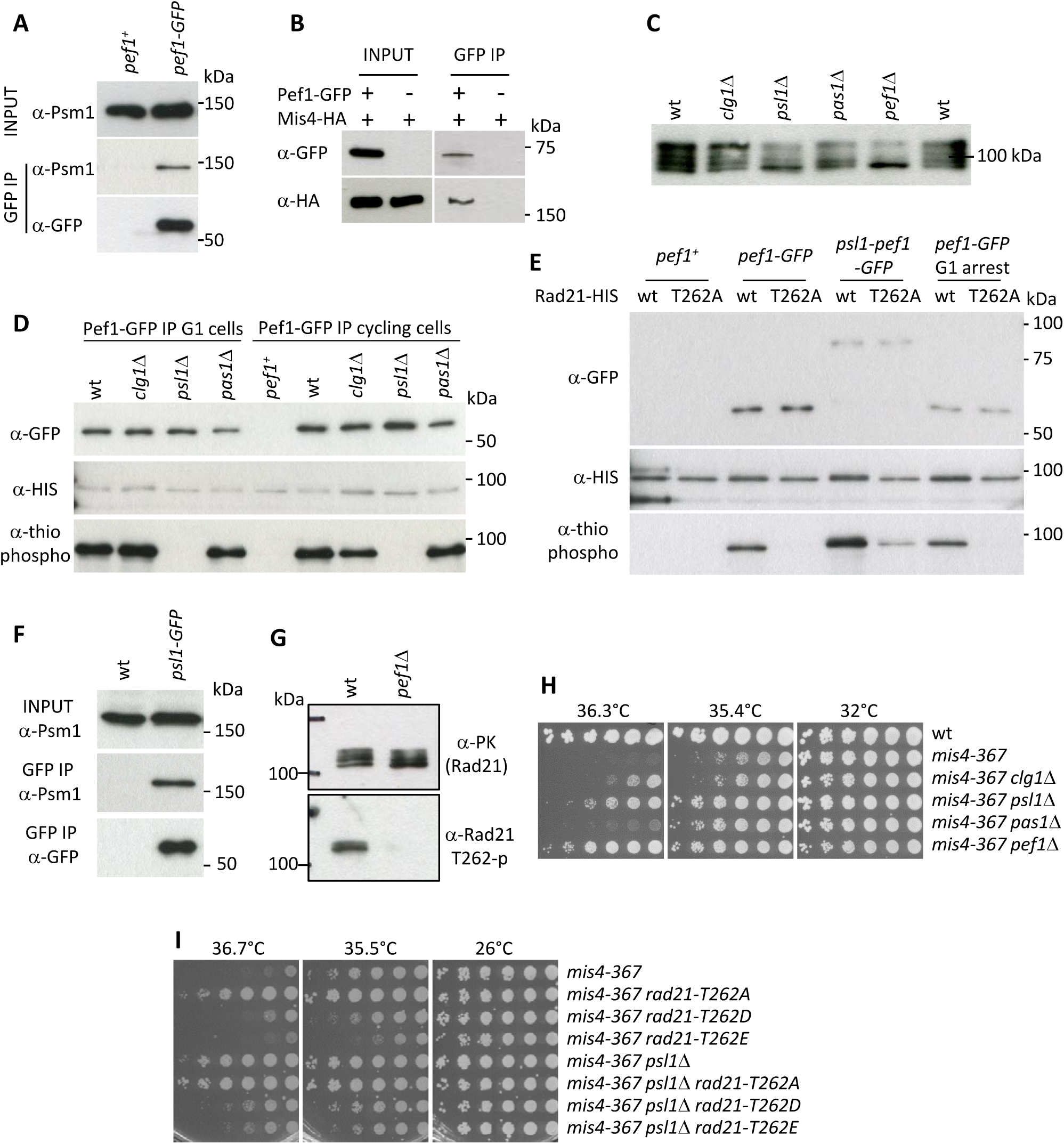
Pef1 phosphorylates Rad21. **(A,B)** Pef1 co-immunoprecipitates cohesin (A) and the cohesin loader Mis4 (B) from total protein extracts. **(C)** Western blot analysis of total protein extracts from cycling cells probed with anti-Rad21 antibodies. **(D)** *In vitro* kinase assays. Pef1-GFP immuno-purified (IP) from cycling or G1 (*cdc10-129*) cells was incubated with *in vitro* translated Rad21-HIS in the presence of ATPγS and the proteins analyzed by Western blotting. Phosphorylated products were detected using an anti-thiophosphate ester antibody. **(E)** *In vitro* kinase assays. Rad21-T262A prevents Rad21 phosphorylation by Pef1. The fusion protein Psl1-Pef1-GFP phosphorylates Rad21. **(F)** Psl1 co-immunoprecipitates Psm1 from total protein extracts (cycling cells). **(G)** Rad21-PK was immuno-purified from cycling cells and probed by western blotting with the indicated antibodies. **(H,I)** Growth assays for suppression of the ts growth defect of *mis4-367*.

The *in vivo* relevance of this pathway was assessed by looking at genetic interactions with the cohesin loader mutant *mis4-367*. Individual deletion of the three cyclin genes indicated that *pas1*Δ was a poor *mis4-367* suppressor; the deletion of *clg1* had a weak effect while *psl1*Δ showed the strongest effect (Fig. 5H). Still, the suppression by *psl1*Δ was weaker than that conferred by *pef1*Δ, suggesting that all three cyclins may act with Pef1 to regulate Mis4 function, likely through the phosphorylation of a set of substrates. Importantly, *rad21-T262A* suppressed the thermosensitive growth defect of *mis4-367* (Fig. 5I). The level of suppression was similar to *psl1*Δ, consistent with Rad21-T262 being the main relevant substrate of Pef1/Psl1. Conversely, the phospho-mimicking allele *rad21-T262E* exacerbated the ts phenotype of *mis4-367* and compromised the suppression by *psl1*Δ.

From these data we conclude that all three known Pef1 CDK complexes contribute to cohesin regulation, suggesting multiple relevant substrates. One of these is Rad21. The Pef1/Psl1 CDK phosphorylates Rad21 on residue T262 and genetic analyses suggest that this phosphorylation event may negatively regulate Mis4 function *in vivo*.

### The phosphorylation status of Rad21-T262 contributes to regulating Mis4 binding to Cohesin Associated Regions

Since Pef1 regulates Mis4 binding to CARs on chromosomes and Rad21 is a substrate of the CDK, we asked whether *rad21-T262A* would recapitulate some aspect of Pef1 loss of function. ChIP analyses in G1 arrested cells indicated that Mis4 binding was indeed increased at some loci in a *rad21-T262A* background although the effect was weaker than for the *pef1* deleted strain (Fig. 6B). The most prominent effects were observed at the telomere site Tel1-R (1.5 fold increase) and within centromeric heterochromatin (1.4 and 1.5 fold at *imr2-L* and *dg2-R*, respectively). By contrast, Mis4 binding was close to wild-type levels along chromosome arm sites and within the rDNA gene cluster (28S and NTS). No additive effect was seen in combination with *pef1*Δ, consistent with Rad21-T262 being a substrate of the CDK. Mis4 binding was marginally reduced in the phospho-mimicking mutant *rad21-T262E* but importantly, increased Mis4 binding in *pef1*Δ was reduced in a *rad21-T262E* background (Fig. 6C).

**Figure 6.**
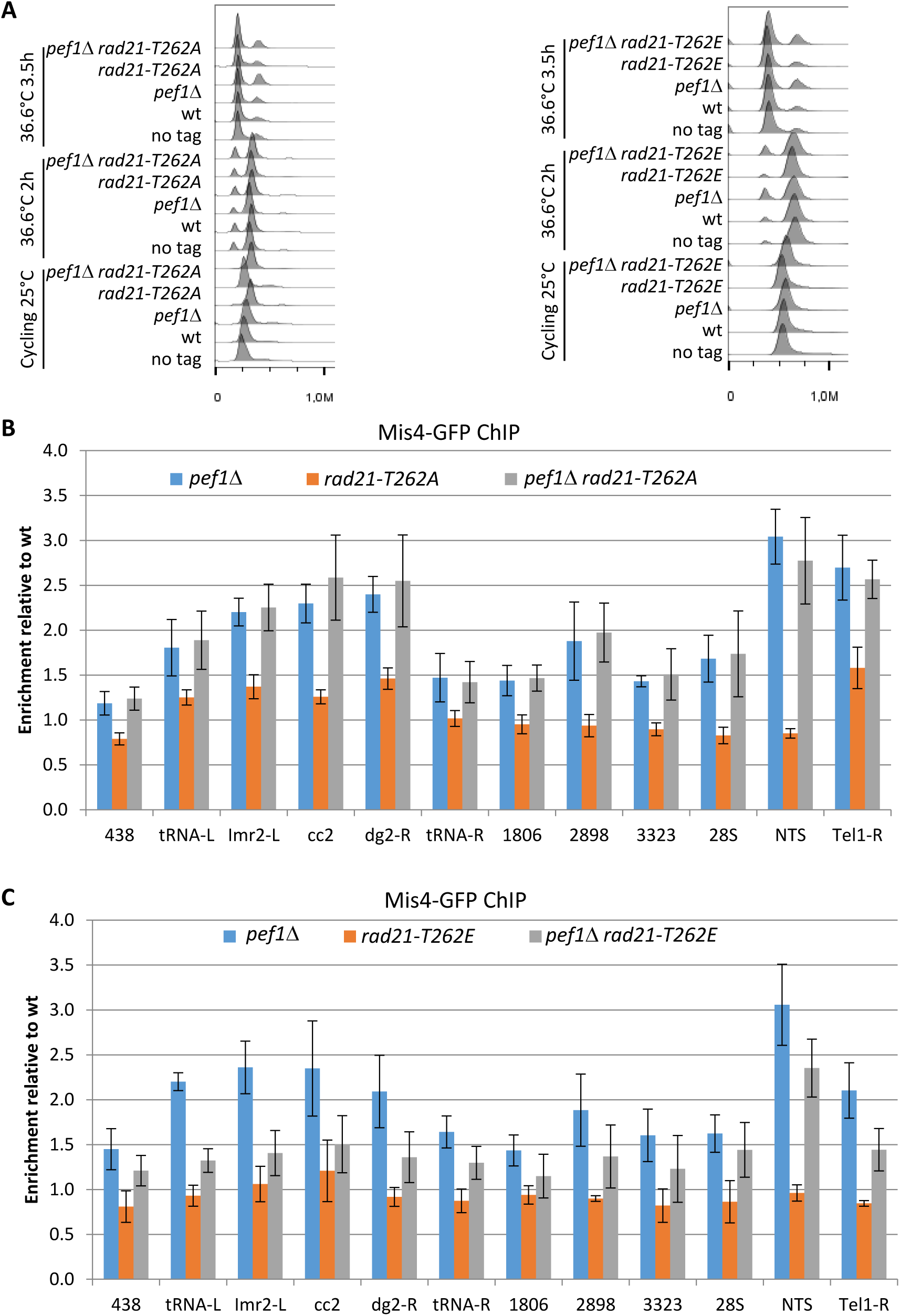
Rad21-T262 phosphorylation modulates Mis4 binding to G1 chromosomes. **(A)** DNA content analysis. Cultures of the indicated strains were shifted to 36.6°C to induce the *cdc10-129* arrest and cells collected for ChIP after 3h. **(B,C)** Mis4-GFP ChIP relative to wild-type. Bars indicate mean ± SD from 4 ratios.

In summary, phosphorylated Rad21-T262 is necessary but not sufficient for Pef1-mediated down-regulation of Mis4 binding. Reciprocally, non-phosphorylatable Rad21-T262 by itself is sufficient to enhance Mis4 binding to specific loci but does not fully recapitulate Pef1 ablation. We conclude that Pef1 regulates Mis4 binding to its chromosomal sites through the phosphorylation of a set of substrates, including Rad21-T262.

### Pef1 and Protein Phosphatase 4 oppose each other

We previously reported that Protein Phosphatase 4 regulated the phosphorylation state of the cohesin subunit Rad21 [40]. Western blot analyses indicate that slow migrating Rad21 isoforms accumulate in a strain deleted for *pph3*, encoding the catalytic subunit of PP4 ([40] and Fig. 7A). Conversely, fast migrating Rad21 isoforms accumulate in a *pef1* deleted strain and a mixed pattern is observed when both the CDK and PP4 are ablated (Fig. 7A), suggesting that Pef1 and PP4 may oppose each other for controlling the phosphorylation status of Rad21.

**Figure 7.**
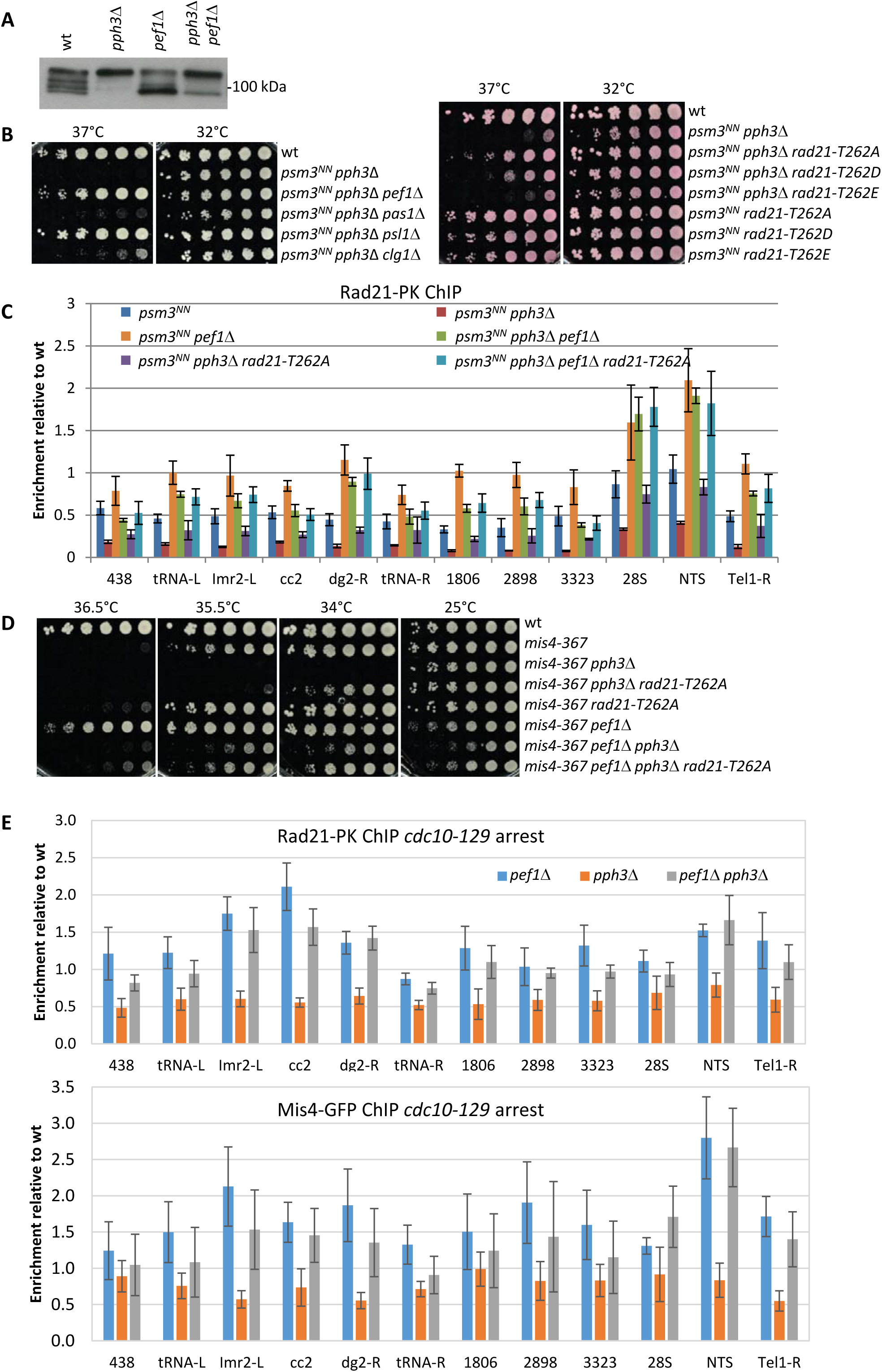
Pef1 and PP4 oppose each other for regulating Rad21 binding to chromosomes. **(A)** Western blot analysis of total protein extracts from cycling cells probed with anti-Rad21 antibodies. **(B)** Cell growth assays indicate that *pef1* and *pph3* show opposite genetic interaction with *psm3^NN^*. **(C)** Rad21-ChIP after one complete cell cycle at 36°C. Data are expressed relative to wild-type. Bars indicate mean ± SD from 4 ratios. **(D)** Cell growth assays showing that *pef1*Δ, *rad21-T262A* and *pph3*Δ display opposite genetic interactions with *mis4-367*. **(E)** Rad21 and Mis4 ChIP relative to wild-type in *cdc10*-*129* arrested cells. Bars indicate mean ± SD from 4 ratios.

A link between PP4 and cohesin loading was provided through the analysis of genetic interactions. Acetyl-mimicking forms of Psm3 are known to inhibit DNA capture by cohesin *in vitro* and reduce the amount of chromatin-bound cohesin *in vivo* [13; 55]. A negative genetic interaction was observed between *pph3*Δ and *psm3^K105NK106N^*(*psm3^NN^*) an acetyl-mimicking allele of *psm3* [35]. The double mutant strain was unable to grow at elevated temperature (Fig. 7B). Interestingly, growth was efficiently rescued by the deletion of *pef1*, *psl1*, and by the non phosphorylatable allele *rad21-T262A*. The phospho-mimicking allele *rad21-T262E* alone did not recapitulate the effect of deleting *pph3*, suggesting that the negative genetic interaction involves the accumulation of other phosphorylated substrates.

ChIP analyses (Fig. 7C) confirmed that Rad21 binding to chromosomes was reduced in a *psm3^NN^*background after one complete cell cycle at 36°C at all sites examined. PP4 ablation exacerbated this phenotype whereas Pef1 ablation has the opposite effect. Therefore in a context of compromised cohesin loading (*psm3^NN^*), PP4 activity stimulated cohesin loading while Pef1 restrained it. Consistent with the genetic data, poor Rad21 binding in the absence of PP4 was efficiently rescued by *pef1*Δ and to a lower extent by *rad21-T262A* (Fig. 7C). It is of note that Rad21 binding was slightly higher in *pef1*Δ than in *pef1*Δ *pph3*Δ at most chromosomal sites, indicating that full stimulation of Rad21 binding by Pef1 ablation requires functional PP4. In the absence of Pef1 some substrates may be phosphorylated by another kinase and de-phosphorylated by PP4. The rDNA gene cluster behaved differently. Rad21 was bound to a similar extent in *pef1* and *pef1 pph3* deleted strains, suggesting that no other kinase phosphorylates Pef1 targets at these loci.

In summary, this experiment revealed that Pef1 and PP4 oppose each other in a situation where the cohesin loading reaction is compromised. Consistently, a similar set of genetic interactions were observed when the activity of the cohesin loader was compromised by the *mis4-367* mutation (Fig. 7D).

To see the effect of PP4 and Pef1 in otherwise wild-type cells, we looked at Rad21 and Mis4 binding to G1 chromosomes by ChIP. PP4 ablation lead to an overall decrease of DNA bound Rad21 and Mis4 (Fig.7E). Conversely, both Rad21 and Mis4 binding were stimulated in the *pef1* deleted strain. The double mutant strain showed a profile similar although not identical to *pef1*Δ alone. This is consistent with Pef1 and PP4 opposing each other for controlling the phosphorylation state of common substrates: the phosphorylated state (*pph3*Δ) reduces cohesin loading while the non-phosphorylated state (*pef1*Δ and *pef1*Δ *pph3*Δ) has the opposite effect. The differences observed between *pef1*Δ and *pef1*Δ *pph3*Δ suggests that other kinase(s) may be contributing.

### Rad21 phosphorylation status may modulate the activity of the cohesin loader

The *psm3^NN^ pph3*Δ strain grew poorly and spontaneous suppressors were frequently observed. Genetic analyses showed that the vast majority were allelic to *pef1*. However, four suppressors were allelic to *mis4* and efficiently rescued *psm3^NN^ pph3*Δ growth and chromosome segregation defects (Fig. 8A, D). Strikingly, all five mutations clustered within the hook domain of Mis4 (Fig. 8B). Even more striking, the very same mutations were recovered as intragenic suppressors of the ts allele *mis4-G1326E* [56] suggesting they enhance Mis4 activity. This region is enriched for residues mutated in Cornelia de Lange syndrome and Kikuchi *et al.* have shown that many of these mutations specifically disrupt the Scc2-Scc1 interaction ([19] and Fig. 8C). In Mis4, this region does not contain any CDK consensus or reported phosphorylation site suggesting it may not be targeted by Pef1/PP4. However, since Rad21 is hyper-phosphorylated in PP4-deprived cells, the suppressor mutations may help Mis4 accommodating a phosphorylated substrate. This possibility is consistent with the finding that *pef1*Δ and *rad21-T262A* mutants are efficient suppressors of *psm3^NN^ pph3*Δ, and that Rad21 phosphorylation was indeed reduced in *pef1*Δ *pph3*Δ when compared to *pph3*Δ alone (Fig. 7A). We suggest that the activity of the cohesin loader may be modulated by the phosphorylation of its cohesin substrate.

**Figure 8.**
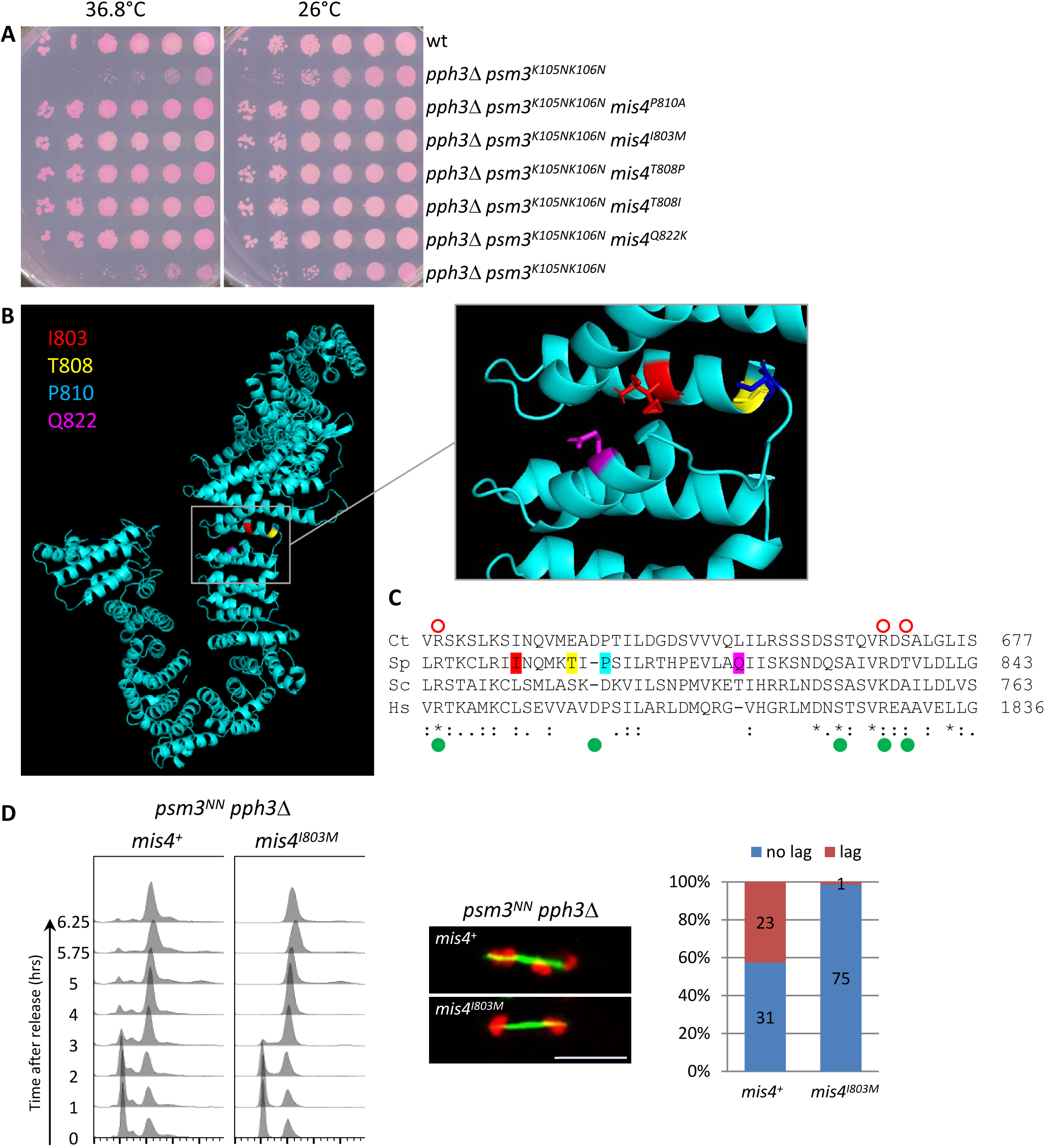
Mis4^I803M^ suppresses *pph3*Δ *psm3^NN^*chromosome segregation defects. **(A)** Cell growth assay. The *mis4* mutations restore *pph3*Δ *psm3^NN^*growth at elevated temperature **(B)** Model structure of Mis4 showing the location of the mutated residues. **(C)** Sequence alignment of Mis4 with Scc2 proteins from other species. The mutated residues in Mis4 are colored as in (B). Ct Scc2 residues required for Scc1 binding are indicated by open red circles and Hs Scc2 residues mutated in Cornelia de Lange syndrome are indicated by green dots as in [19]. Ct, *Chaetomium thermophilum*; Sc, *Saccharomyces cerevisiae*; Sp, *Schizosaccharomyces pombe*, Hs, *Homo sapiens*. **(D)** Mis4^I803M^ efficiently suppressed *pph3*Δ *psm3^NN^* chromosome segregation defects. Cells were arrested in G1 by nitrogen starvation at 25°C and released at 37°C. Progression in the cell cycle was monitored by DNA content analysis. Cells from the 5.75 and 6.25 time points were processed for DNA and tubulin staining to score the number of anaphase cells with lagging chromatids.

## DISCUSSION

Cohesin is involved in a wide range of cellular functions at all stages of the cell cycle, implying a tight control by the cell machinery. The data presented here provides evidence that the CDK Pef1 and PP4 are part of this regulatory network. We will discuss here how phosphorylation may control the activity of the cohesin loader and speculate about the potential physiological implications.

In otherwise wild-type cells Pef1 ablation increased the interaction of both cohesin and its loader Mis4 with their regular binding sites on chromosomes. This concerted increase suggests that cohesin deposition is enhanced because more Mis4 molecules have been recruited to CARs. These cohesin complexes appeared functional as Pef1 inhibition improved sister-chromatid cohesion and chromosome segregation in a *mis4-367* background. Therefore, the most straightforward interpretation is that reduced cohesin loading activity in *mis4-367* is enhanced when the CDK is inactivated, providing sufficient cohesin amenable to sister chromatid cohesion establishment at the time of S phase. Alternatively, cohesin and its loader may accumulate there as the result of a delay in completing some aspect of the loading reaction. Besides DNA capture cohesin form intra-chromosomal loops which may be generated by a distinct biochemical activity of cohesin, possibly regulated by the loading complex [14; 57; 58]. The CDK may positively regulate the formation of loops. In this scenario, the inhibition of the CDK may rescue sister-chromatid cohesion in Mis4 compromised cells by increasing the pool of cohesin available for cohesion at the expense of those engaged in DNA looping. Further studies will address this attractive possibility.

What would be the physiological role for a negative regulator of cohesin loading? As mentioned above, the CDK may regulate non-cohesive aspects of cohesin functions with consequences on gene expression and nuclear architecture. Pef1 ablation in otherwise wild-type cells did not lead to obvious chromosome segregation defects, suggesting that increased cohesin deposition in G1/S may not have any adverse effect on the establishment of sister chromatid cohesion. However, *pef1* mutants showed a negative genetic interaction with *eso1-H17* which is deficient for cohesin acetylation [35], suggesting a positive role for Pef1 in sister chromatid cohesion establishment or maintenance that would remain cryptic when Eso1 is fully functional. A G1/S control of cohesin may also be relevant to DNA damage response during these stages of the cell cycle. In support of these ideas the closest Pef1 human homolog, CDK5, has been implicated in DNA damage response and gene regulation [59].

Understanding how Pef1 regulates cohesin binding to DNA will require further knowledge of the biochemical activities of both cohesin and its loader. Mis4/Ssl3 interacts with all cohesin subunits and these contacts contribute to the activity of the cohesin loader [20]. Structural studies revealed a high conformational flexibility of Scc2 suggesting that the loader may capture cohesin by making multiple contacts around the surface of the ring that may help conformational changes required for DNA capture [18]. Of particular interest for the present study is the interaction between the loader and Rad21. We identified Rad21-T262 as a Pef1 substrate which when phosphorylated contributes to down regulating cohesin binding to CARs. Threonine 262 is located within the central, unstructured domain of Rad21. Interestingly, T262 lies between the two Mis4/Ssl3 contact sites (145–152 and 408–422) that were mapped on Rad21 by peptide arrays [20] and adjacent to the Scc2 binding site (126–230) on *Chaetomium thermophilum* Scc1 [19]. Rad21 phosphorylation may therefore hinder or modify Mis4 interaction with cohesin. Our co-immunoprecipitation assay is consistent with this possibility as the interaction between Mis4 and cohesin appeared slightly increased in the absence of Pef1. Another argument came from the suppressors of the negative interaction between *psm3^NN^* and *pph3*Δ. Besides *pef1* mutants and *rad21-T262A*, suppressor mutations were found in Mis4 and clustered within a Rad21 binding domain [19]. These amino acid changes may help accommodate hyper-phosphorylated Rad21 when PP4 is ablated.

The negative charges bring about by phosphorylated residues may also hinder or modify cohesin interaction with DNA. A recent study indicates that cohesin would tether DNA in its smaller lumen that is, between Rad21 and the SMC’s heads [60]. The two acetylatable lysine residues within Psm3 head domain are thought to stimulate the ATPase activity of cohesin when in contact with DNA [20]. Rad21 phospho-residues may alter the path of the DNA along Rad21 and hinder its contact with Psm3 lysine sensors. Alternatively or additionally, Rad21 phosphorylation may affect the recently reported DNA binding interface between Scc1 and Scc3, the respective budding yeast counterparts of Rad21 and Psc3 [61]. In essence, hypo-phosphorylated cohesin may favor the interaction with either of both the cohesin loader and DNA, and reciprocally, targeted phosphorylation events may have the opposite issue. A phosphorylation based control of cohesin is appealing as these modifications are reversible and occur within seconds with high special resolution. The CDK / PP4 module may provide a fast and accurate mechanism for translating cellular cues into an appropriate cohesin response. We have shown here that Pef1 ablation rescued sister-chromatid cohesion defects of a crippled cohesin loader. Such a regulation may impinge on the other functions of cohesin such as DNA repair and intra-chromosomal looping. Considering the conservation of the Pef1 CDK and PP4 across species, a similar regulation may operate in larger eukaryotes, including humans.

## MATERIALS AND METHODS

### Strains, media and genetic techniques

General fission yeast methods, reagents and media are described in [62]. All strains are listed in Table 1. Experiments were carried out using YES medium unless otherwise stated. Gene deletions and epitope tagging were performed by gene targeting using polymerase chain reaction (PCR) products [63]. The strain carrying an ectopic copy of *rad21-FLAG* was constructed by integrating a *tetO7-rad21-FLAG* construct into a gene free region on chromosome 3 [64]. The tetracycline sensitive repressor was introduced by crossing with *ura4^+^-tet^ON^*(*tetR-tup11D70* integrated at the *ura4* locus [65]). Expression was induced by the addition of 5µg/ml tetracycline (anydrotetracycline hydrochloride, SIGMA, stock solution 10mg/ml in DMSO) or DMSO alone for the un-induced control.

**TABLE 1.**
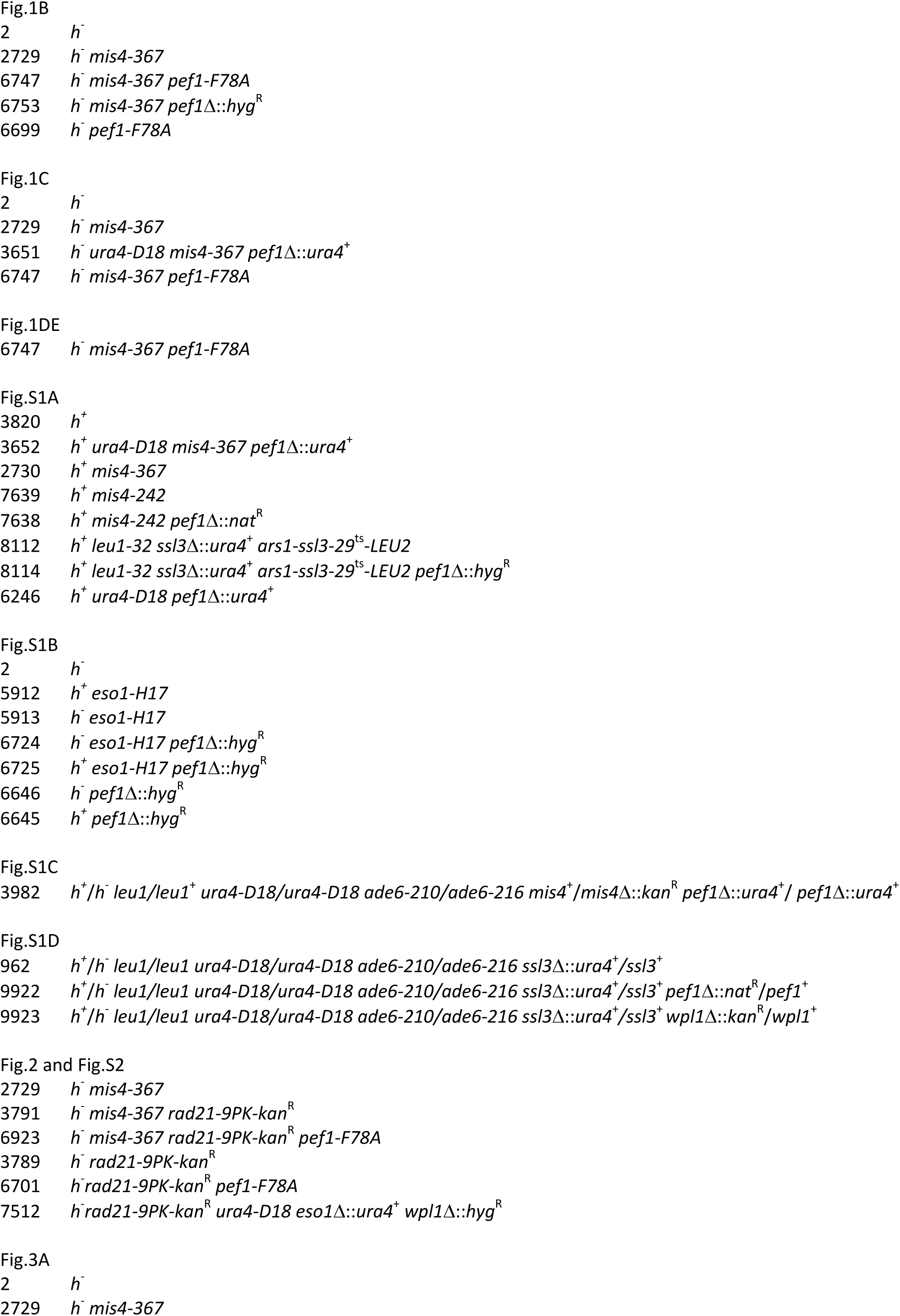
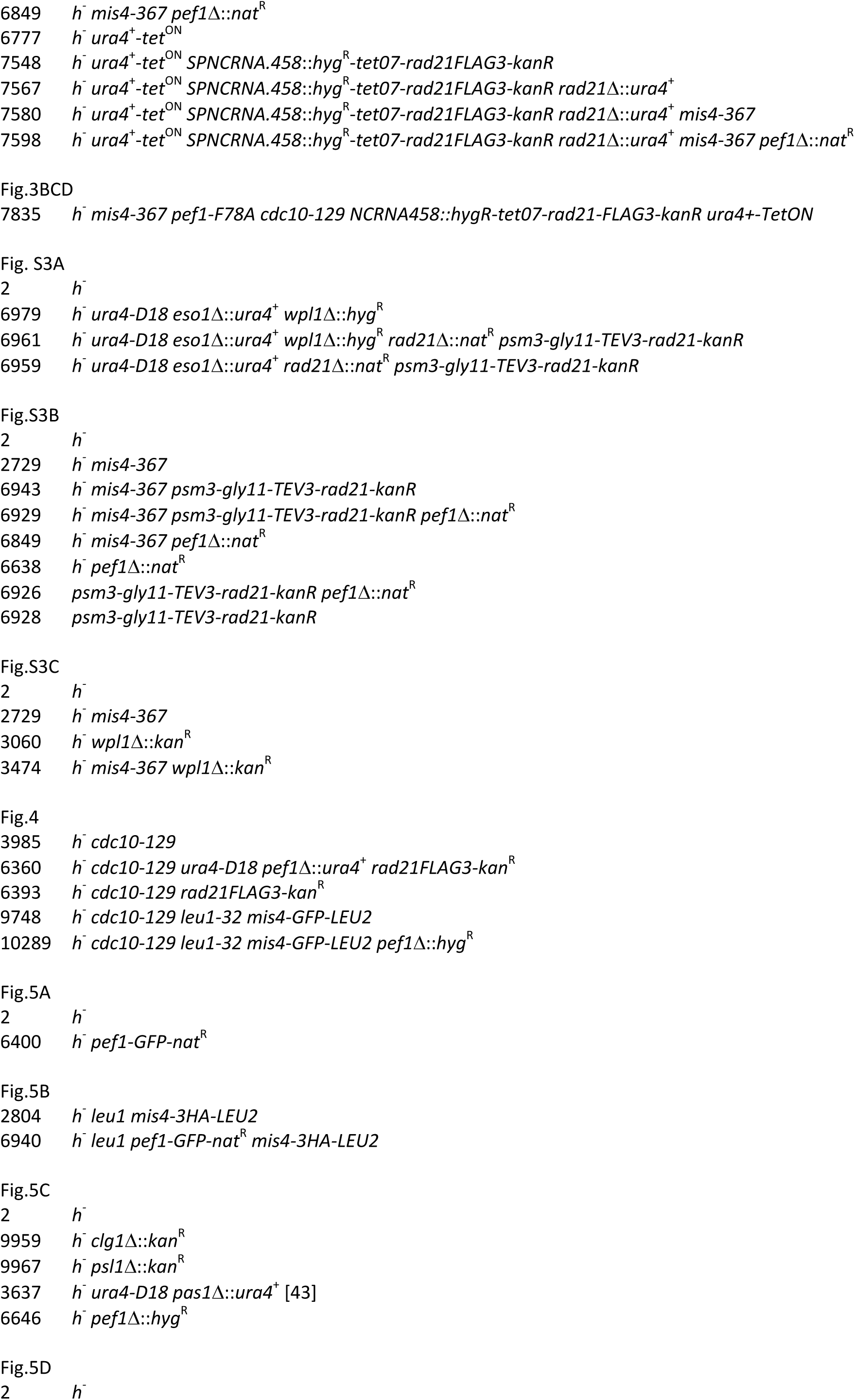
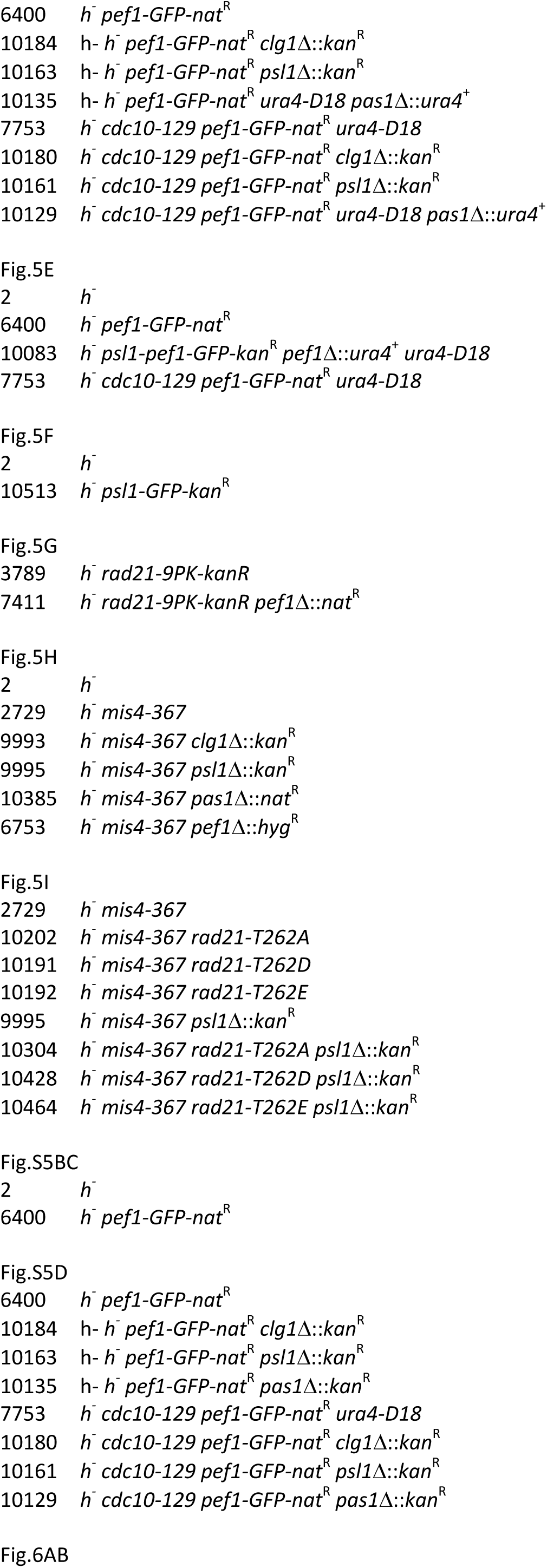
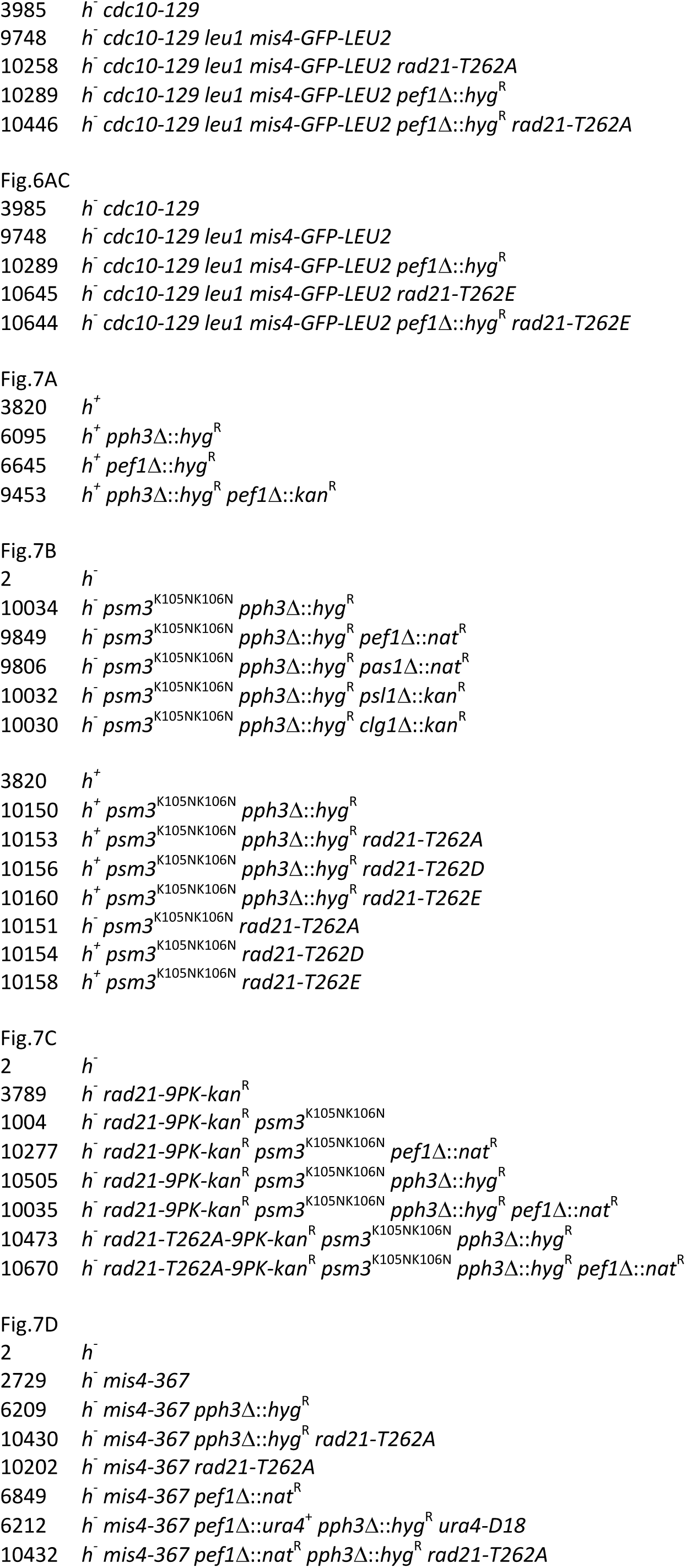
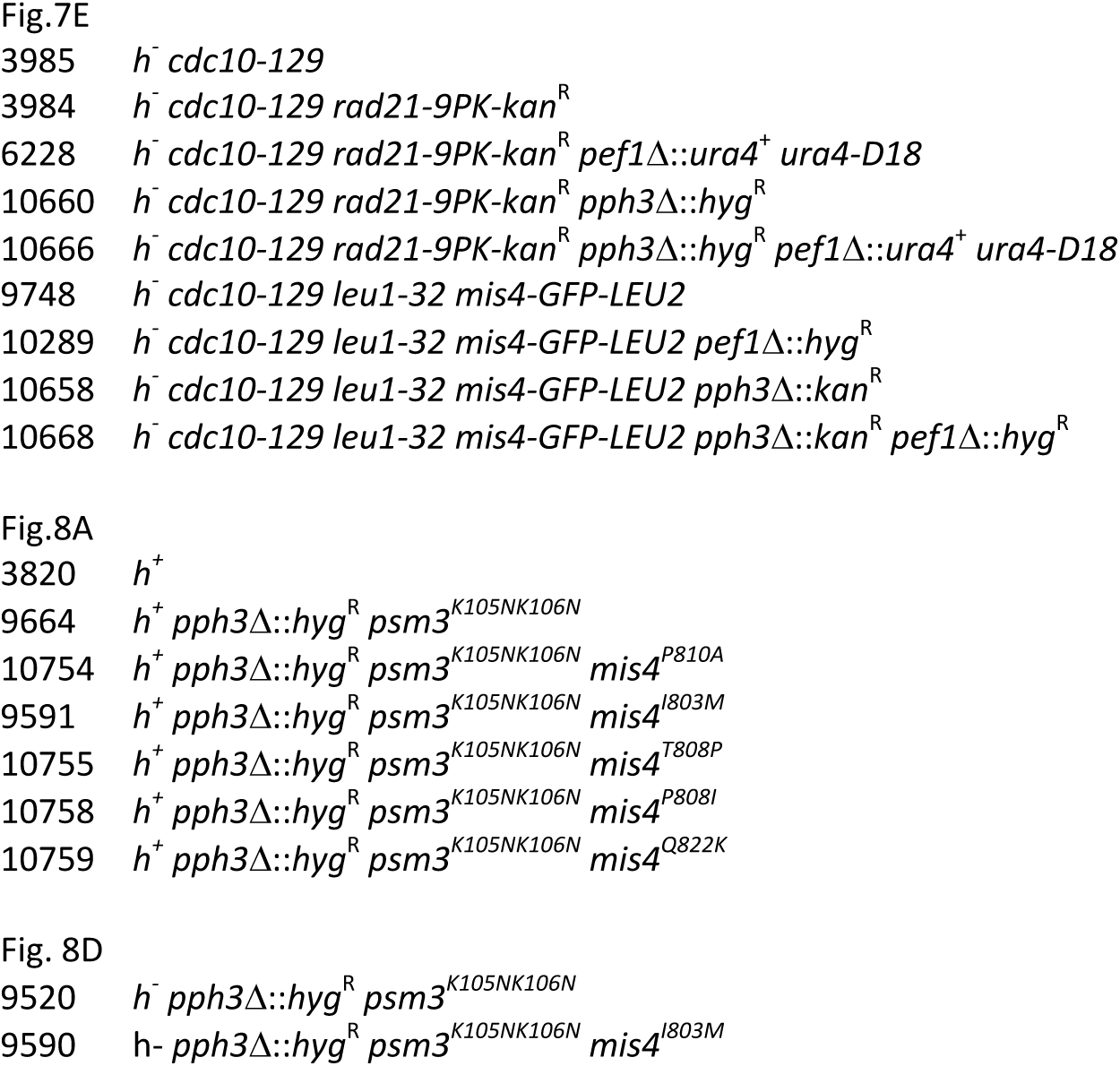
Strain list.

Suppressors of the *mis4-367* thermosensitive growth phenotype were obtained either with or without UV mutagenesis. Cells were plated onto YES medium, irradiated with UV light to ∼50% killing, and incubated at 25°C until colony formation. For spontaneous suppressors, the UV irradiation step was omitted. Colonies were replica plated onto YES plates containing the vital dye Phloxin B and incubated over night at 37°C. Suppressors appeared as white growing colonies in an otherwise background of red-stained dead cells. Suppressors were backcrossed at least three times. Eleven suppressors showed a monogenic segregation and fell into 4 linkage groups. Genetic mapping of group 1 mutants indicated linkage with the *cds1* and *sds21* loci on chromosome III. The mutated locus was identified by Comparative Genome Hybridization (CGH). Genomic DNA was extracted from the mutant strain *sup^UV12^* and the wild-type *S. pombe* reference strain (SP972) and co-hybridized to a CGH tiling array (29-32 mer probes with 7 or 8 base spacing from the start of one probe to the start of the next, Roche Nimblegen). The array spanned 1,290,000 bp of chromosome 3, from coordinates 117000 to 1065000 and from 1143000 to 1485000 (Genbank NC_003421.2 GI:63054406). DNA regions carrying candidate Single Nucleotide Polymorphisms (SNPs) were used to design a high resolution tiling array (29-30 mer probes tiled such as each candidate SNP is analyzed by 8 probes, 4 on each DNA strand). A single A to G SNP (N146S) was found within SPCC16C4.11 (*pef1*). The mutation was confirmed by PCR and DNA sequencing. The *pef1* gene was deleted and genetic analyses showed that *pef1*Δ was allelic to *sup^UV12^* and suppressed the Ts phenotype of *mis4-367*.

The *pef1-as* allele (*pef1-F78A*) was generated by *in vitro* mutagenesis. The DNA fragment carrying the mutated allele was transformed into a recipient strain in which the region of interest was substituted by the *ura4^+^* marker (*pef1d(260-32)::ura4*^+^), and Ura^−^ colonies were selected on 5-Fluoroorotic acid plates. The *pef1-as* allele in the selected strain was amplified by PCR and checked by sequencing. The inhibitor 1-NA-PP1 (Cayman Chemical, stock solution 25mM in DMSO) was added to the culture medium at 25μM. The equivalent volume of solvent alone (DMSO) was added for the control sample. The construction of *rad21* alleles was done using the strategy described in [40].

### Cytological techniques

DNA content was measured by flow cytometry with an Accuri C6 Flow cytometer after Sytox Green staining of ethanol-fixed cells [66]. Data were presented using the FlowJo software. Immunofluorescence and Fluorescence In Situ Hybridization (FISH) were done as described [67]. Briefly, cells were fixed by the addition of paraformaldehyde to a final concentration of 1.8% in 1.2M sorbitol. The flasks were removed from 36°C, incubated at 21°C for 45 min and processed for tubulin staining using TAT1 antibodies [68]. Cells were re-fixed and processed for FISH using the centromere linked c1228 cosmid as a probe [69]. Cells were imaged using a Leica DMRXA microscope and a 100X objective. Tubulin staining was used to select cells with an interphase array of microtubules. Distances between FISH signals were measured from maximum projections of images created from z series of eight 0.4-μm steps using MetaMorph software. Cen2FISH signals were considered as separated when the distance was greater than 0.3µm. Statistical analysis was done using two-tailed Fisher exact test with 95% confidence interval. Nuclear spreads were done as described [35]. Signal intensity was measured in a square surface containing the spread nucleus. Background signal was measured by moving the square surface to an adjacent region devoid of nuclei. The background value was subtracted for each nucleus. The signal was quantified for at least 35 nuclei for each sample. The mean and the confidence interval of the mean were calculated with α = 0.05.

### Antibodies, protein extracts, immunoprecipitation, Western blotting, cell fractionation, chromatin immunoprecipitation (ChIP) and kinase assay

Rabbit polyclonal antibodies against Rad21, Psm1, Psm3, Psm3-K106Ac have been described previously [35; 70]. The mouse monoclonal anti-tubulin antibody TAT1 is from [68]. Anti-Rad21-T262P antibodies were raised by Biotem (Apprieu, France). Rabbits were immunized with the KLH-coupled peptide C+SVTHFSTpPSMLP. Sera were immune-depleted by affinity with the non-phosphorylated form of the peptide and antibodies were affinity purified against the phosphorylated peptide. Other antibodies were of commercial source. Rabbit polyclonal anti-GFP A11122 (Molecular Probes), mouse monoclonal anti-GFP (Roche), anti-PK antibodies (monoclonal mouse anti V5 tag, AbD serotec), mouse monoclonal anti-FLAG (Sigma), rabbit polyclonal anti-histone H3 (Abcam 1791) and rabbit monoclonal anti-thioester (Abcam 92570).

Protein extracts, immunoprecipitation (IP), cell fractionation and western blotting were as described [35; 48]. Chromatin Immunoprecipitation (ChIP) was as described in [40] using anti-FLAG, anti-PK or anti-GFP (A11122) antibodies. ChIP enrichments were calculated as percentage of DNA immunoprecipitated at the locus of interest relative to the input sample. The mean was calculated from 4 technical replicates with error bars representing standard deviation. Enrichments relative to wild-type were calculated as the mean of 4 ratios with errors bars representing standard deviation.

For kinase assays, the Rad21-6HIS substrates were produced using a coupled transcription/translation reaction system (*E. coli* EasyXpress, biotechrabbit) using plasmid DNA templates. The reaction was carried out at 37°C for 1h using 5–10 nM plasmid DNA in a total volume of 50µL. Pef1-GFP was immunoprecipitated (IPed) from total cell extracts (5×10^8^cells) prepared in lysis buffer (50mM Hepes pH 7.6; 75mM KCl; 1mM MgCl2; 1mM EGTA; 0.1% Triton X-100; 1mM DTT; 10mM Sodium butyrate; Glycerol 10%) supplemented with inhibitors (Protease inhibitor cocktail Sigma P8215, 1mM PMSF, 1mM Na vanadate, 20mM β-glycerophosphate). The IPed material was washed three times with 0.2ml of lysing buffer without inhibitors, and twice with 1.5X kinase buffer (75mM TRIS pH 7.5; 15mM MgCl2; 1.5mM EGTA; 1.5mM DTT). The CDK bound to the beads was recovered in 60µL of the 1.5X kinase buffer. The kinase assay was done using 40µL of CDK beads and 20µL of *in vitro* generated Rad21-6HIS. The reaction was carried out with 1mM ATPγS at 37°C for 1h with 1min shaking (300rpm) every 10min. The reaction was stopped by adding EDTA to 20mM. To alkylate thio-phosphorylated proteins p-nitrobenzyl mesylate (Abcam 138910) was added to 5mM and the samples incubated for 2 hours at 21°C on a rotating wheel. The beads were removed by loading the sample on a magnetic column equilibrated with kinase buffer. The flow-through was collected and the presence of thio-phosphorylated proteins was assayed by Western blotting with anti-thioester antibodies (1/10000 dilution).

## AKNOWLEDGMENTS

We thank our colleagues M.-F. Giraud and S. Manon for their initial help and advice for the *in vitro* production of proteins and kinase assays, K. Gull for the gift of anti-tubulin antibodies, K. Tanaka and H. Okayama from providing *pef1* and *pas1* deleted strains. This work was supported by the Centre National de la Recherche Scientifique, l’Université de Bordeaux, la Région Aquitaine, l’Association pour la Recherche sur le Cancer (PJA 2013 1200 205; PJA 20171206211) and l’Agence Nationale de la Recherche (ANR-14-CE10-0020-01). Adrien Birot was supported by a fellowship from the Agence Nationale de la Recherche Investissements d’Avenir ANR-10-IDEX-03-02 and l’Association pour la Recherche sur le Cancer (DOC20160603884). Amélie Feytout was supported by a fellowship from the Ministère de l’Enseignement Supérieur et de la Recherche. Karl Ekwall was supported by grants from the Swedish Cancer Society (CF) and the Swedish Research Council (VR).

## COMPETING INTEREST

The authors declare no competing interests.

---

**Figure 1 supplement.**
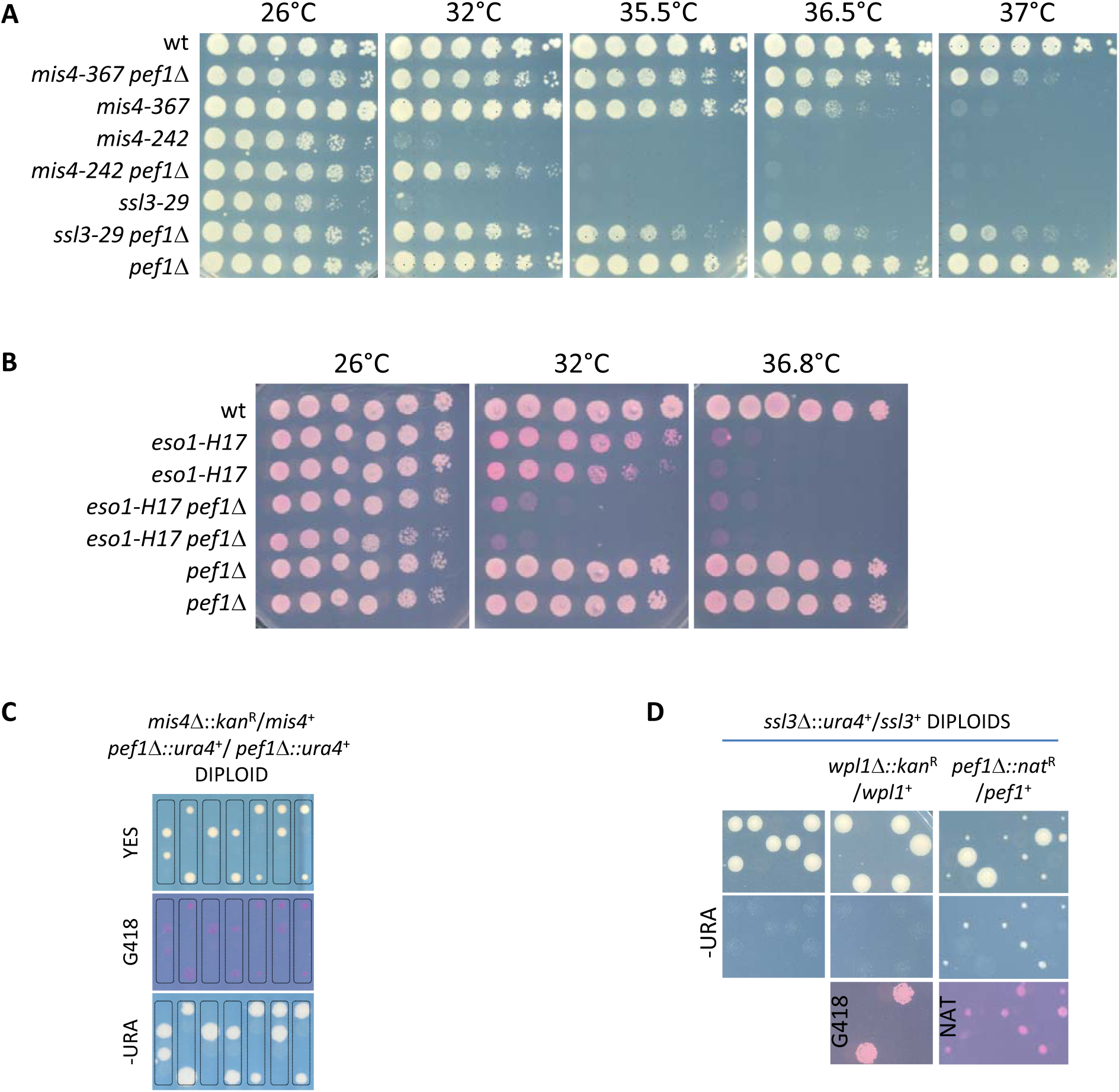
**(A)** Deletion of the *pef1* gene alleviates the growth defects of cohesin loader mutants. **(B)** Pef1 ablation exacerbates the thermosensitive growth phenotype of the cohesin acetyl-transferase mutant *eso1-H17*. **(C)** Deletion of the *pef1* gene does not bypass Mis4 requirement. Tetrad dissection from a *mis4*Δ::kanR/*mis4*^+^ *pef1*Δ*::ura4*^+^/ *pef1*Δ*::ura4*^+^diploid strain. Among the 4 spores of a tetrad, only two form viable colonies. All viable colonies are *mis4*^+^. (D) Deletion of *pef1* allows cell survival in the absence of the *ssl3* gene. The *wpl1* or *pef1* gene was deleted in a *ssl3*Δ::*ura4*^+^/*ssl3*^+^ diploid strain. The resulting diploids were sporulated an tetrads dissected. In an otherwise wild-type (left), or *wpl1*Δ/*wpl1*^+^ (middle) background, Ura^+^ (*ssl3*Δ) progeny was not observed. By contrast small growing Ura+ NatR (*ssl3*Δ *pef1*Δ) colonies were frequent among the progeny from the *pef1*Δ/*pef1*^+^ diploid (right).

**Figure 2 supplement.**
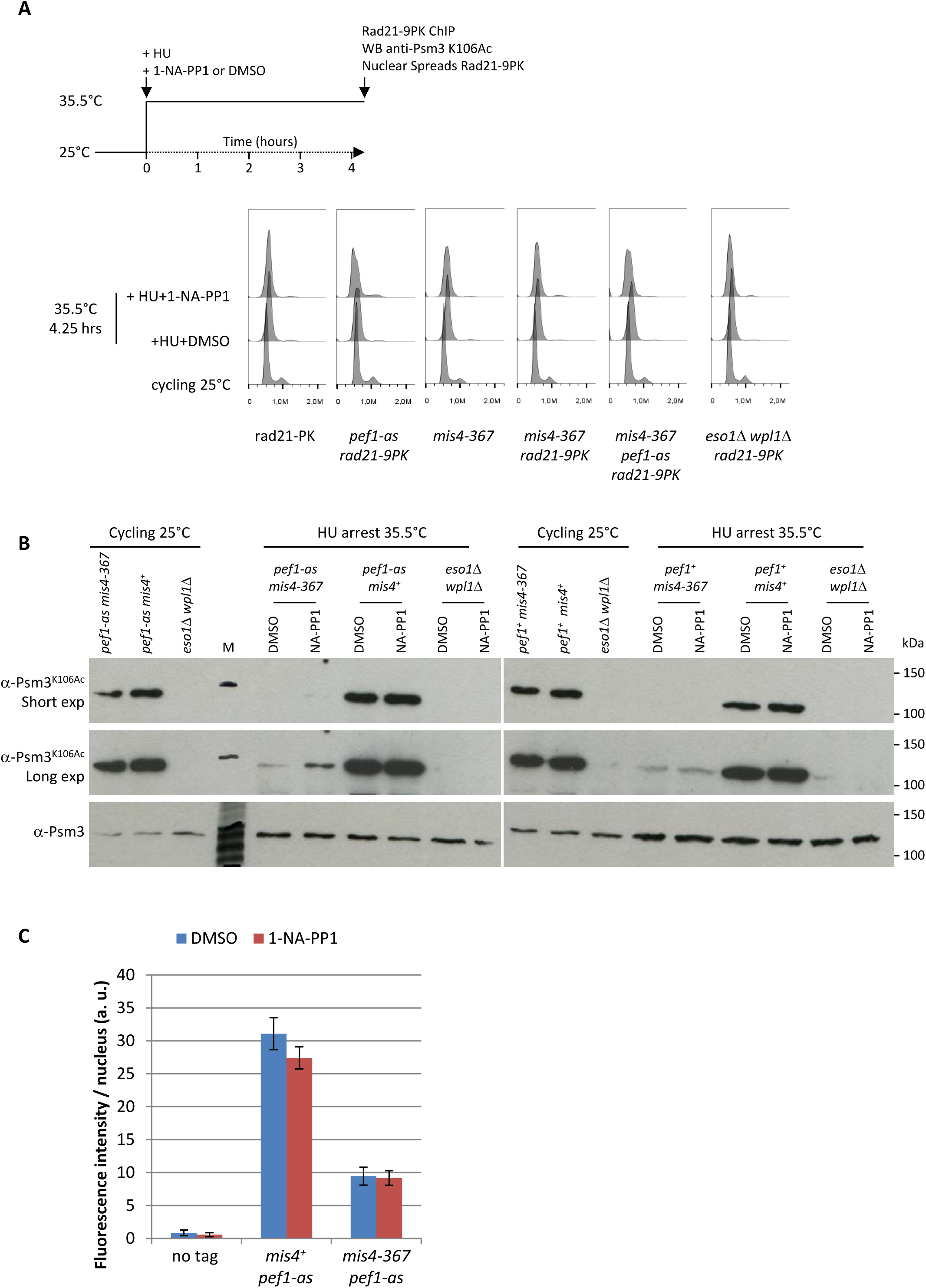
**(A)** DNA content analyses. The drift in the 1C peak to the right in HU-arrested cells is due to an increase in the mitochondrial DNA content as the cells elongate [71]. **(B)** Inhibition of Pef1 kinase activity increased the level of Psm3 acetylation in *mis4-367* cells at the restrictive temperature. Total protein extracts were analyzed by western blotting using the indicated antibodies. M: molecular weight markers. **(C)** The inhibition of Pef1 kinase activity did not induce a global change in the total amount of chromatin-bound Rad21. The amount of chromatin-bound Rad21-9PK per nucleus was measured by nuclear spreads and indirect immunofluorescence using anti-PK antibodies. The bars represent the mean fluorescence intensity per nucleus +/- the 95% confidence interval.

**Figure 3 supplement.**
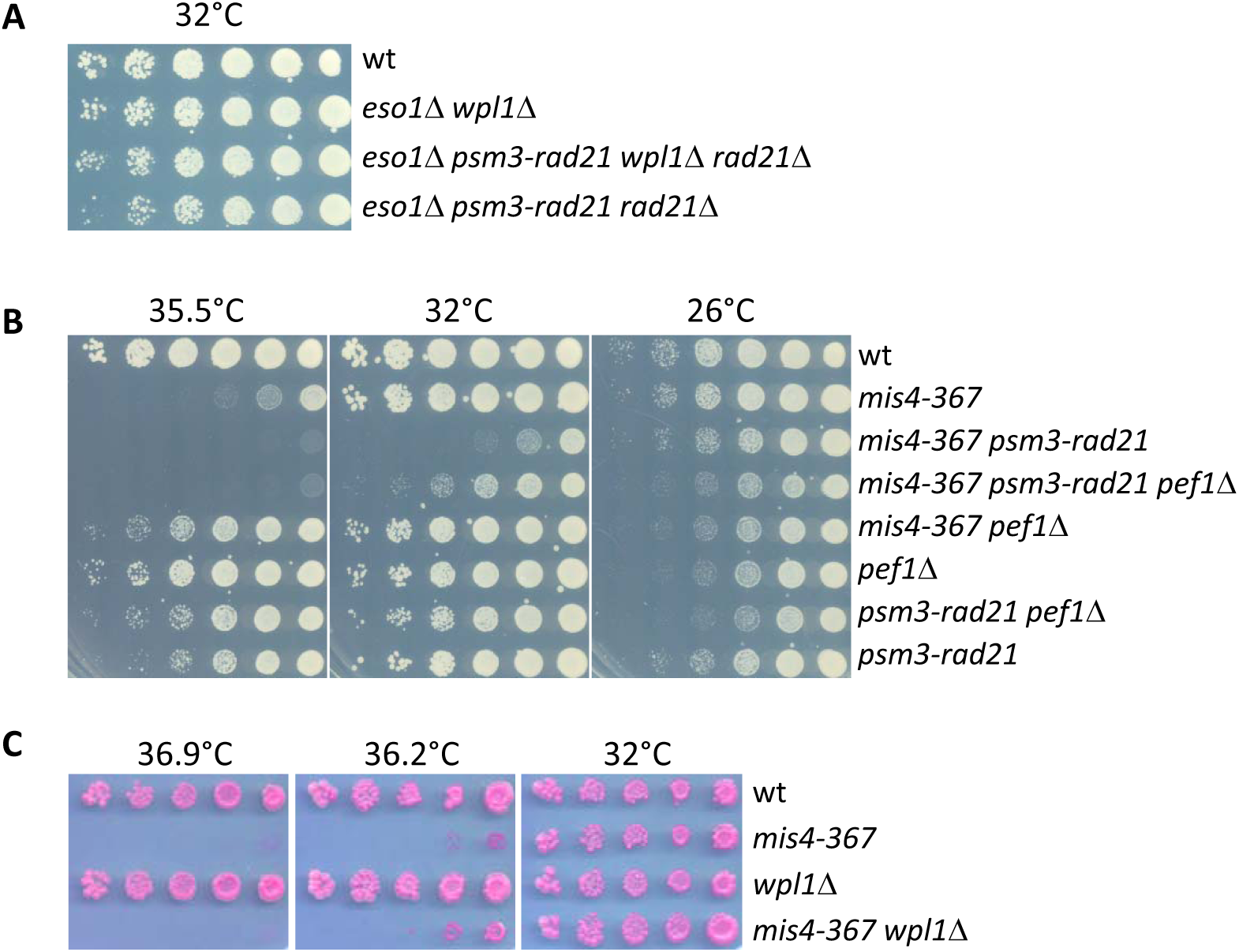
Pef1 acts independently from the Psm3/Rad21 interface. **(A)** Deletion of *wpl1* or a *psm3-rad21* gene fusion allows cell survival in the absence of the otherwise essential *eso1* gene. **(B)** The *psm3-rad21* gene fusion does not suppress the *mis4-367* thermosensitive growth defect. However deletion of the *pef1* gene improves growth in this genetic setup. **(C)** Deletion of the *wpl1* gene does not suppress *mis4-367* growth defect.

**Figure 5 supplement.**
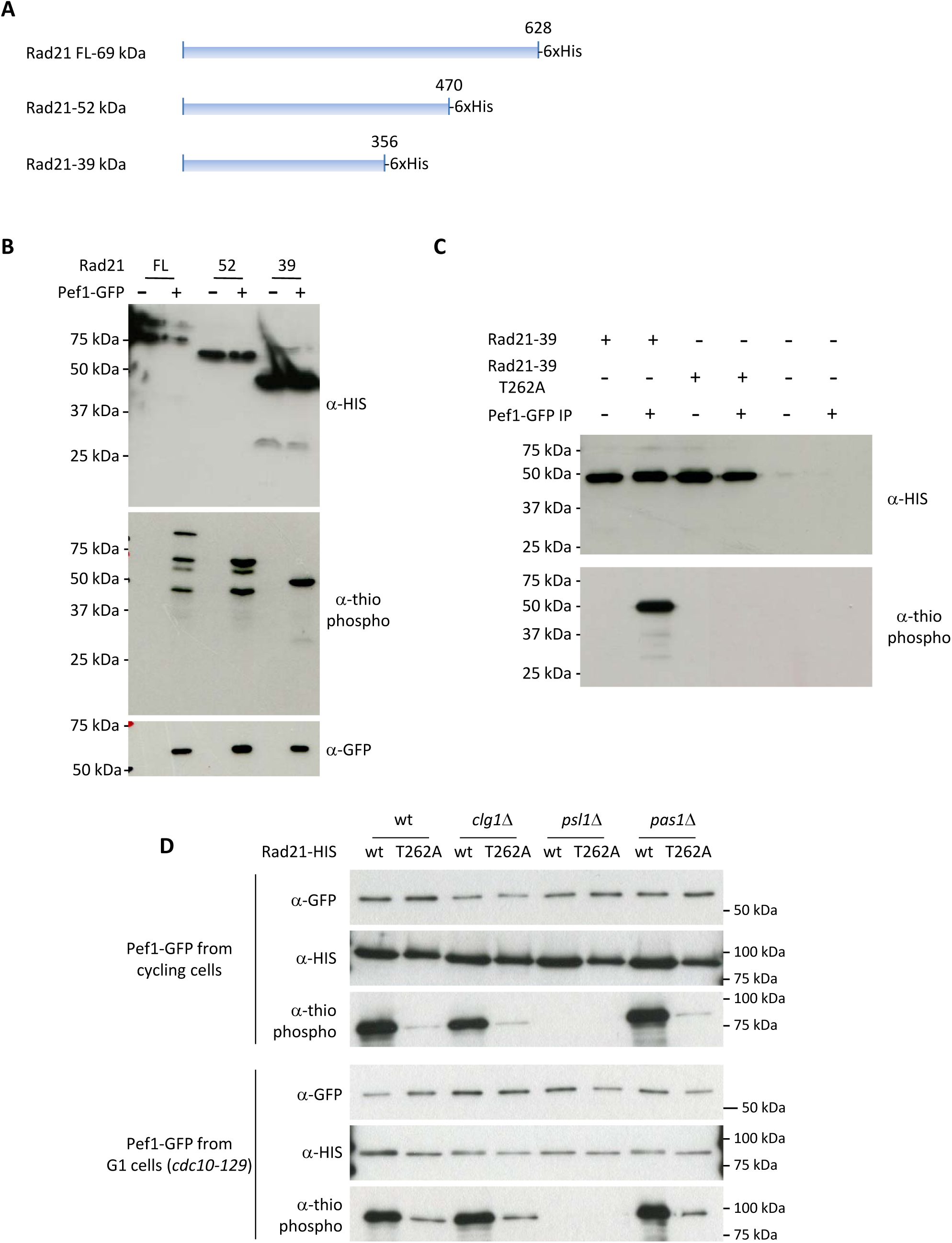
Mapping the Pef1 phosphorylation site within Rad21. **(A)** Truncated forms of Rad21 used in the kinase assays. **(B-C)** *In vitro* kinase assays using Pef1-GFP purified from cycling cells. The reaction products were analyzed by western blotting with the indicated antibodies. Note that all Rad21 forms migrate slower than predicted from their calculated molecular weight. The shortest Rad21 derivative (Rad21-39) is efficiently phosphorylated by Pef1. The T262A substitution abrogates *in vitro* phosphorylation of Rad21-39 by Pef1. **(D)** Full length Rad21-T262A is a poor Pef1 substrate.

